# A disease similarity approach identifies short-lived Niemann-Pick type C disease mice with accelerated brain aging as a novel mouse model for Alzheimer’s disease and aging research

**DOI:** 10.1101/2024.04.19.590328

**Authors:** Vikas Anil Gujjala, Isaiah Klimek, Morteza Abyadeh, Alexander Tyshkovskiy, Naci Oz, José Pedro Castro, Vadim N. Gladyshev, Jason Newton, Alaattin Kaya

**Affiliations:** Department of Biology, Virginia Commonwealth University, Richmond, VA 23284 USA; Center for Integrative Life Sciences, Virginia Commonwealth University, Richmond, VA 23284 USA; Division of Genetics, Department of Medicine, Brigham and Women’s Hospital, Harvard Medical School, Boston, MA 02115, USA; i3S, Instituto de Investigação e Inovação em Saúde, Universidade do Porto, 4200-135, Porto, Portugal; Aging and Aneuploidy Laboratory, IBMC, Instituto de Biologia Molecular e Celular, Universidade do Porto, 4200-135, Porto, Portugal; Department of Human and Molecular Genetics, Virginia Commonwealth University, Richmond, VA, 23284, USA

**Keywords:** Comparative Genomics, Niemann Pick Disease Type C, Alzheimer’s Disease, Congenital Diseases, Biomarkers

## Abstract

Since its first description in 1906 by Dr. Alois Alzheimer, Alzheimer’s disease (AD) has been the most common type of dementia. Initially thought to be caused by age-associated accumulation of plaques, in recent years, research has increasingly associated AD with lysosomal storage and metabolic disorders, and the explanation of its pathogenesis has shifted from amyloid and tau accumulation to oxidative stress and impaired lipid and glucose metabolism aggravated by hypoxic conditions. However, the underlying mechanisms linking those cellular processes and conditions to disease progression have yet to be defined. Here, we applied a disease similarity approach to identify unknown molecular targets of AD by using transcriptomic data from congenital diseases known to increase AD risk, namely Down Syndrome, Niemann Pick Disease Type C (NPC), and Mucopolysaccharidoses I. We uncovered common pathways, hub genes, and miRNAs across *in vitro* and *in vivo* models of these diseases as potential molecular targets for neuroprotection and amelioration of AD pathology, many of which have never been associated with AD. We then investigated common molecular alterations in brain samples from an NPC disease mouse model by juxtaposing them with brain samples of both human and mouse models of AD. Detailed phenotypic and molecular analyses revealed that the NPC^mut^ mouse model can serve as a potential short-lived *in vivo* model for AD research and for understanding molecular factors affecting brain aging. This research represents the first comprehensive approach to congenital disease association with neurodegeneration and a new perspective on AD research while highlighting shortcomings and lack of correlation in diverse *in vitro* models. Considering the lack of an AD mouse model that recapitulates the physiological hallmarks of brain aging, the characterization of a short-lived NPC mouse model will further accelerate the research in these fields and offer a unique model for understanding the molecular mechanisms of AD from a perspective of accelerated brain aging.

## Introduction

Alzheimer’s disease (AD) is the most common dementia caused by frontotemporal neurodegeneration. It progresses with age, and current therapeutic options are limited to mitigating symptoms rather than treating the underlying disease(1). Despite copious research and effort into the disease, no biomarkers are properly characterized for AD causation and progression. With the lack of accurate biomarkers for early identification and proper medical interventions to prevent or treat the disease, AD has become the most prevalent form of dementia in the elderly(2). Although initial research suggests that extracellular amyloid plaques and intracellular tau tangles are common pathologies for AD, recent findings point to a more compelling cause: lysosomal storage disorders and energy homeostasis disruption, both of which are associated with congenital conditions characterized by dementia-like pathology(3). Interestingly, these cellular pathologies are also the hallmark of several congenital disorders with neurodegenerative pathologies that are also associated with increased risk for AD. Among them, discussed here are Down syndrome (DS), Niemann Pick Disease Type C (NPC), and Mucopolysaccharidosis I (MPS I). Recent studies have also shown that more than 50% of Down Syndrome patients suffer from early-onset AD(4). MPS I is known to cause severe cognitive decline and neuronal loss(5), leading to early-onset AD-like pathology.

Down Syndrome, also referred to as Trisomy 21, resulting from supernumerary chromosome 21, is one of the most frequent survivable aneuploidies. The triplication of genes on chromosome 21 leads to several abnormalities, ranging from musculoskeletal disorders and congenital heart disease to neurodegeneration and cognitive decline. DS patients are more susceptible to autoimmune conditions, hematological disorders, cognitive decline, and inverse comorbidity with solid-state tumors, all pathologies associated with AD. DS increases the risk of early-onset AD due to accelerated aging and cognitive decline(6,7).

Niemann Pick Disease Type C is a rare autosomal progressive lysosomal storage disorder resulting from *NPC1* and/or *NPC2* gene mutations. The mutations in *NPC1* and *NPC2* genes render them dysfunctional, resulting in the accumulation of lipids in various tissues due to the inability to internalize cholesterol and other lipids. This accumulation, in turn, leads to cognitive decline as well as neurological and psychiatric disorders. The survival rate of NPC at birth is meager, and the number of patients crossing infancy is minimal, making it one of the rarest congenital disorders ever studied, limiting the data available. NPC has an elevated risk of developing severe early-onset AD(8). NPC is also referred to as childhood AD and leads to the development of the most severe AD pathology among those discussed here and uniquely uncouples AD from aging.

Mucopolysaccharidoses are a set of inherited metabolic disorders characterized by the dysfunction or absence of one or more enzymes required for processing glucosaminoglycans, leading to their accumulation and homeostasis disruption in various tissues, including the brain, spinal cord, and nervous system. MPS I is also known to increase the risk of developing early-onset AD(9).

Although studies have been investigating these disorders and their potential risk for AD individually (10–13), there have been no reports applying the disease similarity approach on a molecular level to identify common, potentially targetable similarities for early disease diagnosis and developing drug candidates against the pathological onsets of AD. Accordingly, we constructed a disease-to-disease similarity network using transcriptomics data across different sample types, including frontal cortex brain samples, brain organoids, and iNSCs (*induced neural stem cells*) for each disease. We then assessed connections and common molecular risk factors between AD and DS, NPC, and MPS I. We identified previously unknown genes, and pathways that contribute the most to the similarities among the disorders, and identified regulatory gene networks and novel miRNAs as molecular targets. We further extended our comparative genomics approach by comparing frontal cortex transcriptome data from a transgenic *NPC1* mutant (*Npc1^tm(I1061T)Dso^*) mouse model which expresses the most common human mutation found in NPC patients(14) to the AD mouse model(15) and compared their similarities to postmortem human brain samples. Along with phenotypic and chronological and biological aging data, our findings from the disease similarity approach suggest that the NPC1^mut^ mouse model can be utilized as a unique, short-lived *in vivo* model for understanding AD mechanisms and identified molecular factors that contribute to brain aging and AD-like pathogenesis. Overall, our data advances a new perspective on AD research while highlighting shortcomings and lack of correlation in diverse *in vitro* models, which can accelerate diagnostic and therapeutic applications against common types of dementia.

## Materials and Methods

### Data acquisition and analysis

The raw count data for various sample types and diseases were collected from NCBI GEO (https://www.ncbi.nlm.nih.gov/geo/) when the following conditions were satisfied; (i) the data collected matches the sample type required, and (ii) inclusion of proper control samples.

We collected or extracted the raw transcriptomics data from GEO from the raw matrix files using Seurat and SingleCellExperiment packages in R. The data was filtered by removing repeats, missing values or blanks, and expression values without valid gene names and then passed to trimmed mean (TMM) normalization to harmonize the data. We performed Differential expression analysis with R package DESeq2(16). We declared gene expression to be significantly changed if fold change (FC) was 1.5 or higher in any direction (log2FC = 0.585) and False Discovery Rate (FDR) less than 0.05 for human genes and only imposed the FDR less than 0.05 without constraints on fold change for mouse genes. We generated the Volcano plots using the ggplot2 package in RStudio. The TMM normalized data were used for heat map generation using RNAlysis software(17).

We conducted a thorough commonality analysis on the differentially expressed genes identified from prior analyses, aiming to identify the common genes across different diseases. We validated these genes against the random commonness expected from DEGs using Fisher’s exact test for randomness, ensuring a comprehensive assessment of the false discovery rates for non-randomness as applicable for each sample type.

### Venn Diagrams

We generated the Venn diagrams to visualize common DEGs found earlier from various diseases or sample types using InteractiVenn software (http://www.interactivenn.net/#) and processed them using Inkscape (https://inkscape.org/). The outputs generated were text files containing the data of the genes overlapping between the data sets and the Scalable Vector Graphics (.svg) files(18).

### Gene Network Construction

The Protein-Protein interaction and protein-microRNA interaction networks were generated by the Network Analyst and processed using Cytoscape(19,20).

We specified the “organism” as *H. sapiens*, the “ID type” as the Official gene symbol, and a list of genes in a sequence to Network Analyst to generate various interaction networks. The Analysis overview page provided the options: Protein-protein interaction (PPI), gene regulatory network (GRN), Diseases, and Gene Coexpression Networks.

The brain-specific gene-miRNA interaction networks were constructed from the miRNet database (https://www.mirnet.ca/miRNet/home.xhtml) using the list of differentially expressed genes as the query. Gene-miRNA interactions referred to the miRTarBase v8.0 database as the standard. In Cytoscape, we used a discrete mapping option to alter the shape, color, and font size of specific nodes in the network. The degree of interaction/association between nodes in the network was used as the score to make the color spectrum, and the nodes were colored based on their respective scores.

### Enrichment plots and Heat maps

Functional enrichment analyses were performed by using EnrichR(21) and the outcomes were visualized by using ShinyGO(22). The heat map was generated based on the TMM normalized raw data, and it had the following specifications: downregulated genes represented in blue, upregulated genes in red, and bidirectional complete clustering done with the Euclidean distance method. Each disease was grouped along with its control. All the groups contained TMM-normalized triplicate RNA-seq data.

### Identifying human orthologs of mouse genes

Two different sources and methods were employed to obtain the human orthologs for the mouse genes obtained from both our RNA sequencing and downstream transcriptomic analysis of NPC1^mut^ mouse model frontal cortex samples and the DEGs from APP/PS1 AD mouse model transcriptomic data collected from NCBI GEO, which are similar to human transcriptomic data.

The first method involved generating the list of human orthologs using the function “orthologs” inherent to the R package “babelgene.” The second method uses the software “Mouse to Human BMD, Fracture, and OA Results” (https://danielevanslab.shinyapps.io/alliston_mouse2human/)(23–25).

### Animal Studies

Animal Studies were conducted in conjunction with the Transgenic Knock-Out Mouse Shared Resource Core at Virginia Commonwealth University (VCU). WT (C57BL/6) and *Npc1^tm(I1061T)Dso^* mutant mice (Strain #027704, The Jackson Laboratory, Bar Harbor, ME, USA)(14) were bred and housed under a protocol approved by the VCU Institutional Animal Care and Use Committee, which has received accreditation from the Association for Assessment and Accreditation of Laboratory Animal Care. All animals were maintained on a 12-hour light/dark cycle and provided food and water *ad libitum*. Frontal cortex samples were dissected from littermate male and female mice brains immediately after cervical dislocation, snap-frozen in liquid nitrogen, and stored at -80°C until analyzed. Mice were monitored daily to identify new litters, determine their lifespan, and weighed weekly beginning at 4 weeks of age. Survival was determined in a small subset of mice (N=9), after which a humane endpoint (105 days) was established to prevent suffering due to the inability to access food or water.

### Motor function assessment

The development of an overt neurological phenotype was monitored by daily visual observation of tremor activity in the animals. The weights of mice were assessed weekly. Cages containing animals identified with neurological symptoms were supplemented with Diet Gel 76A (72-01-5022X) and HydroGel (70-01-5022) from ClearH2O. Several established phenotypical tests were performed to examine the balance and motor coordination of the NPC1^mut^ mice(26). All animals were trained on each apparatus used at 4 weeks of age. Tests were conducted weekly and in sequence, and the data presented represents the last time point (12 weeks) at which the entire cohort could humanely perform all tests. The first test performed was the vertical screen test, where the mouse is placed on the upper edge of a metal screen with 2 mm wires spaced 1 cm apart (25 cm tall x 22 cm wide). Once the animal gripped the screen with all four paws, the screen was lifted vertically about 50 cm above an 8 cm foam mat so that the animal faced the floor. Test completion was scored when the animal turned upward and climbed to the top of the screen within a 60-second time limit. After performing the vertical screen test, the animal was immediately placed at the center of a 2 cm in diameter round wooden beam measuring 120 cm in length with a wooden box at each end. The apparatus was suspended 50 cm above an 8 cm foam mat. Completion of the test was scored when the animal was able to travel 60 cm in either direction to reach the wooden box without falling from the beam within 180 seconds. Once the balance beam test was finished, the animal was immediately transferred to the center of a 35 cm metal bar with a diameter of 3 mm suspended 25 cm above an 8 cm foam mat, allowing it to hang from the bar by the front paws. Completion of this test was scored as the animal’s ability to actively escape to the end of the bar within 30 seconds. Finally, following the coat hanger test and a ten-minute rest period, the animal was placed on the Rota-rod (Maze Engineers, Skokie, IL, USA) to determine the animal’s ability to maintain balance on a 3 cm motor-driven drum rotating at 15 revolutions per minute. Animals were evaluated on their retention time and considered to have completed the test after maintaining active climbing for 360 seconds. A two-way ANOVA was used to assess the statistical significance of the results.

### tAge Prediction and Aging Signature Association Analysis

To assess the transcriptomic age (tAge) of brains derived from control mice and NPC1^mut^ mice, we applied a Bayesian Ridge mouse multi-tissue gene expression clock of relative chronological age, lifespan-adjusted age, and expected lifespan based on previously identified signatures of aging and lifespan-regulating interventions (27,28). Genes with less than 10 reads in more than 80% of samples were filtered out, resulting in 15,023 expressed genes according to Entrez annotation. Filtered data was then passed to Relative Log Expression (RLE) normalization (29). Normalized gene expression profiles were log-transformed and scaled. The missing values corresponding to clock genes not detected in the data were imputed with the precalculated average values. Samples from control animals were used as the reference group. Differences between average tAges across the groups were assessed with the mixed-effect model using the rma.uni function from the meta for package in R.

The association of changes induced by NPC1^mut^ in the brain with established transcriptomic signatures of aging and hazard was examined at the level of enriched pathways. The utilized signatures included tissue-specific liver, kidney, and brain signatures of aging in mice and rats as well as multi-tissue signatures of aging and expected hazard, unadjusted and adjusted for chronological age(28). For the identification of enriched functions perturbed by NPC1 KO in mouse brains, we first performed differential expression analysis, comparing control and NPC1^mut^ brain samples separately for each sex as well as across sexes including sex as a covariate, using edgeR. We then performed functional GSEA(30) on a pre-ranked list of genes based on log10(p-value) corrected by the sign of regulation, calculated as: -log(pv)×sgn(lfc), where pv and lfc are p-value and logFC of a certain gene, respectively, and sgn is the signum function (equal to 1, -1 and 0 if the value is positive, negative or equal to 0, respectively). HALLMARK, KEGG, and REACTOME ontologies from the Molecular Signature Database (MSigDB) were used as gene sets for GSEA. The GSEA algorithm was performed via the fgsea package in R with 10,000 permutations and a multilevel splitting Monte Carlo approach. p-values were adjusted with the Benjamini-Hochberg method. An adjusted p-value cutoff of 0.1 was used to select statistically significant functions. A similar analysis was performed for gene expression signatures of aging and hazard. Signatures of enriched pathways were compared based on the Spearman correlation of their normalized enrichment scores (NES).

## Results

### Cross-comparison of commonly altered genes between AD and other congenital diseases across different sample types revealed key genes and regulators

Using transcriptomic data from commonly utilized *in vitro* and *in vivo* AD models, including postmortem brain samples, brain organoids, and iPSC-derived neural stem cells (iNSCs) and *in vitro* and *in vivo* models for four different congenital neurodegenerative disorders, we performed comparative genomics-based similarity approaches to identify the common molecular signatures and shared regulators among them (**Figure 1A**). While for AD and DS, we could find data for all sample types, for the rare diseases NPC and MPSI we could only find one sample type for each: brain organoids and iNSCs, respectively (**Supplementary Tables 1 and 2**).

**Figure 1:**
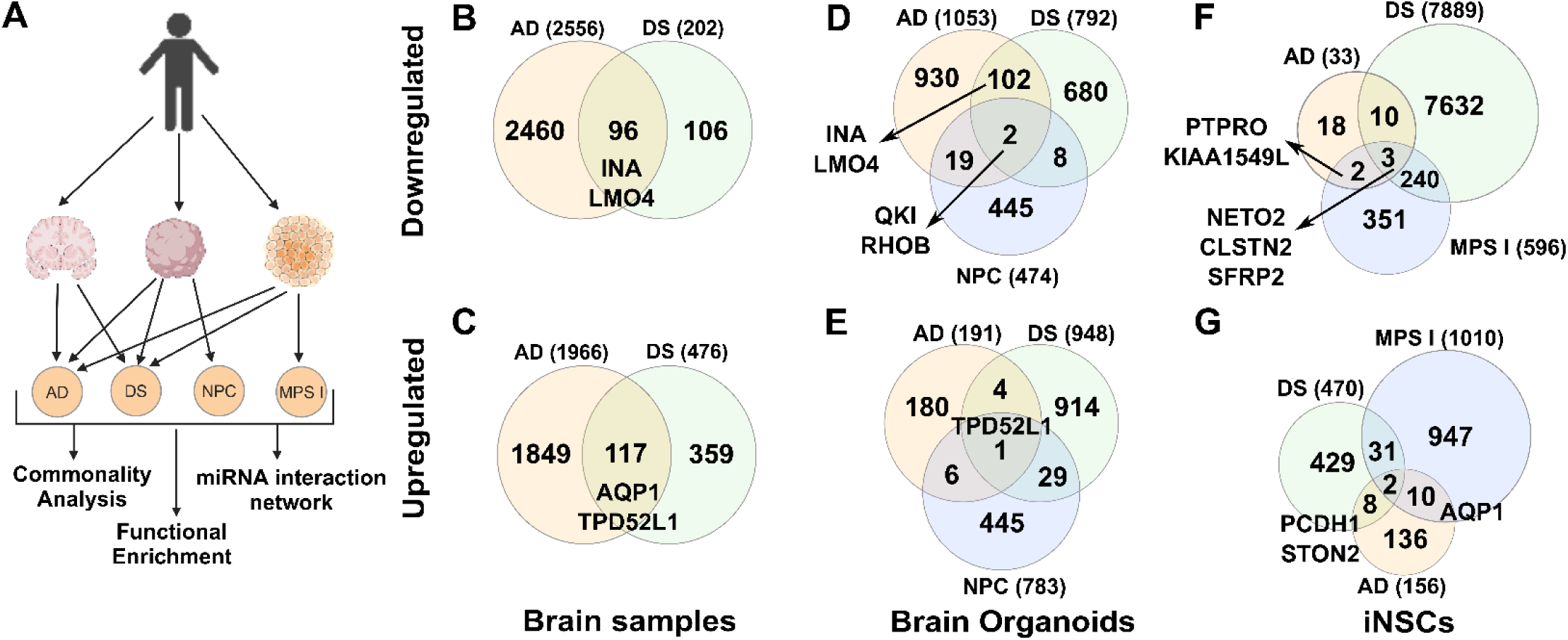
Disease commonality analyses revealed commonly altered genes across different models. **(A)** A summarized description of the disease *in vitro* and *in vivo* models (sample types) was utilized to compare transcriptome data obtained from each sample type and perform downstream analyses across these diseases. These models include frontal cortex Alzheimer’s disease (AD) brain samples, induced neuron-derived organoids, and iPSC-derived neural stem cells (iNSCs). (**B – G)** Venn Diagrams depicting the differentially expressed genes and their commonalities in brain samples (AD and Down Syndrome (DS)), in brain organoids (Neimann-Pick disease type C2 (NPC), AD, DS), and in iNSCs (AD, DS, and Mucopolysaccharidosis type I (MPS I)). The cutoff is 0.585 for log2FC and 1.3 for -log10 (FDR). (**B – E)** There are 2,556 downregulated and 1,966 upregulated genes in AD, 202 downregulated, and 476 upregulated genes in DS frontal cortex brain samples. Among them, 96 genes are downregulated, and 117 are upregulated commonly. (**C – F)** There are 1,053 downregulated and 191 upregulated genes in AD, 792 downregulated and 948 upregulated genes in DS, and 474 downregulated and 783 upregulated genes in NPC organoid samples. Among them, 102 downregulated and 4 upregulated genes are commonly altered between AD and DS. There are 19 downregulated and 6 upregulated genes common between NPC and AD. Two downregulated (*QKI, RHOB*) and one upregulated (*TPD52L1*) genes are common to all three disorders. (**E – G)** In iNSCs, there are 33 downregulated genes in AD, 7,889 in DS, and 596 in MPS I. There are 156 upregulated genes in AD, 470 in DS, and 1010 in MPS I. There are three downregulated and two upregulated genes common across all three disorders.

There were 2,556 downregulated and 1,966 upregulated genes in AD, and 202 downregulated and 476 upregulated genes in Down Syndrome frontal cortex brain samples. Of these, 96 downregulated and 117 upregulated genes significantly overlapped between these two diseases (Fisher’s exact test, *P* = 6×10^-09^) (**Figure 1B, C and Supplementary Table 1**)(31,32). We further analyzed Gene Ontology (GO) enrichment terms for these commonly altered genes. We found that downregulated genes were significantly enriched in pathways associated with GABAergic synapse, Neuroactive ligand-receptor interaction, Glutamatergic synapse, and Oxytocin signaling pathways (**Supplementary Figure 1A**). Interestingly, most upregulated genes were significantly enriched in pathways associated with infectious diseases such as Shigellosis, Toxoplasmosis, Leishmaniasis, Pertussis, and *Staphylococcus aureus* infection. Some of the other pathways were PI3K-Akt signaling, Glutamatergic synapse, and Regulation of the actin cytoskeleton (**Supplementary Figure 1B**).

Next, we compared the expression profiles of brain organoid samples of AD, DS, and NPC(33,34). Brain organoids, generated from induced pluripotent stem cells, hold significant potential by modeling tissue-like environments and recapitulating features of neurodegeneration, as an *in vitro* model(35). The transcriptome data from the NPC organoid samples were the only human-related datasets we could obtain from publicly available data (36). To visualize gene expression variation between organoid samples of these diseases, we performed principal component analysis (PCA). The PCA of the transcriptome data revealed a pattern, with the first three principal components (PCs explaining ∼73 % of the total variance in gene expression with replicates of each disease clustered together among them. While AD and NPC were mainly segregated by PC1, AD and DS samples were segregated by PC2 (**Supplementary Figure 2A**).

To understand the basis of this segregation pattern, we performed a pathway enrichment analysis by combining the 100 top genes (50 with positive weights and 50 with negative weights), contributing to each PC1 and PC2, respectively. Analyses of the genes contributing to PC1 revealed a distinct set of GO terms, such as embryonic hemopoiesis, cytoskeletal organization, generation of neurons and neurogenesis, and cellular protein localization (**Supplementary Figure 2B, Supplementary File 1**). The analysis of genes for PC2 also revealed the overlapping and unique GO terms related to the mitotic cell cycle, head and brain development, viral process, and positive regulation of growth (**Supplementary Figure 2C, Supplementary File 1**). These results suggest that these processes diverged significantly across these diseases and may account for disease progression.

Then, we analyzed gene expression commonalities between these three diseases. We found 131 downregulated and 40 upregulated genes in common, at least among the two diseases (**Figure 1D, E)**. Consistent with the PCA analyses, DS and AD showed significantly (Fisher exact test, *P* = 4.99e-6) overlapped genes that they shared 109 genes commonly altered in between them (104 downregulated, 5 upregulated). In total, 28 genes (21 downregulated, 7 upregulated) were in common between AD and NPC, and 40 genes (10 downregulated, 30 upregulated) were commonly altered between DS and NPC organoid samples. We found only two genes, *QKI* (Quaker homolog, KH Domain RNA binding) and *RHOB* (Ras Homolog Family Member B), downregulated and one gene, *TPD52L1* (Tumor Protein D52 Like 1), upregulated in all three diseases (**Figure 1D, E** and **Supplementary Table 3**).

Finally, the analyses of data from iNSCs also revealed commonly regulated genes across AD, DS, and MPS I (37–39). It should be highlighted here that iNSCs showed diverse transcriptomic profiles for AD and DS. For example, while AD human brain samples have 4582 up- and downregulated genes, the AD model of iNSCs revealed only 186 differentially expressed genes (DEGs) in total. Conversely, iNSCs for the DS model were characterized with 8355 DEGs, while DS brain samples displayed only 688 DEGs. Among them, three genes, Neuropilin and Tolloid Like 2 (*NETO2*), Secreted Frizzled related protein 2 (*SFRP2*), and calsyntenin 2 (*CLSTN2*), were commonly downregulated, and two genes, Growth/Differentiation Factor-15 (*GDF15*), and Adrenomedullin 2 (*ADM2*) were commonly upregulated in AD, DS, and MPS I. Two downregulated genes, Protein Tyrosine Phosphatase Receptor Type O (*PTPRO*) and KIAA1549 Like (*KIAA1549L*), and ten upregulated genes, one of which was Aquaporin 1 (*AQP1*), were common to AD and MPS I (**Figure 1F, G**).

Additionally, considering the observation of diverse transcriptomic changes across different models of AD, we compared the gene expression changes among these experimental models (brain samples, brain organoids, and iNSCs) to identify the common molecular signatures of AD. Application of PCA analyses revealed substantial intra- and intersample heterogeneity (**Supplementary Figure 1A**). We found that the brain samples and organoids had the highest number of commonly altered genes by having 210 downregulated (**Supplementary Figure 3A**) and 22 upregulated (**Supplementary Figure 3B**) genes. Among them, *INA* and *LMO4* genes were commonly downregulated, and *TPD52L1* was among the upregulated genes. It should be noted here that our analyses also identified these genes in commonly altered gene sets of organoid and brain samples of AD, DS, and NPC (detailed in the next chapter) (**Figure 1**). There were 20 genes commonly altered between brain samples and iNSCs. Among them, nine genes, two of which are *PTPRO* (Protein Tyrosine Phosphatase Receptor Type O) and *SPON2* (Spondin 2), are commonly upregulated and 11 of them, including *AQP1* (Aquaporin 1), *PCDH1* (Protocadherin 1), *STON2* (Stonin 1), and *GDF15* (Growth Differentiation Factor 15) were commonly downregulated. Our comparative analyses of AD samples revealed only one gene, *NETO2* (Neuropilin and Tolloid-like 2), commonly downregulated in brain samples, brain organoids, and iNSCs. This data concludes that none of the *in vitro* sample types can fully recapitulate the molecular signatures of Alzheimer’s brain.

The three genes, *INA* (Internexin neuronal intermediate filament protein alpha), *LMO4* (LIM Domain only 4-TF), *TPD52L1* (Tumor protein D52 Like 1), and *AQP1* (Aquaporin 1) were commonly altered across different sample types and different diseases. For example, while *INA* and *LMO4* were commonly downregulated in brain samples and organoids of AD and DS, *TPD52L1* was upregulated in brain samples and organoids across AD, DS, and NPC. The other candidate gene, *AQP1,* was commonly upregulated in brain samples and iNSCs across AD, DS, and MPSI (**Figure 1**).

The *LMO4* gene codes for a transcription factor, with recent research uncovering its association with AD, albeit with decreased expression of *LMO4* correlated with the number of neurofibrillary tangles (NFT) and severity of senile plaque deposition(40). *INA* is crucial for neuronal structure integrity(41). Although overexpression of *INA* has been found to cause abnormal neurofilament accumulations and motor coordination deficits in transgenic mice(42) and α-Internexin aggregates have been associated with neuronal intermediate filament inclusion disease, there has been no report of altered *INA* expression in AD(43,44).

*TPD52L1* is expressed primarily in the cerebral cortex, mainly in glial cells, and in high levels in the reproductive system (45). However, there is a glaring lack of research into this protein and its family of proteins, with most of the focus on *TPD52* and *TPD54* instead. Still, consistent with our data, recent studies suggest slight but significant upregulation of *TPD52L1* in neurodegenerative diseases like Alzheimer’s and Parkinson’s (46,47), but its role and broader impact on the pathology are still relatively unknown. Despite its discovery in 1996 and the characterization of this family shortly after, the functions and mechanisms of the tumor protein D52 family genes, in particular, *TPD52L1*, also referred to as *TPD53*, are relatively understudied, even to this day. While most of the research on this family is focused on *TPD52* and more recently *TPD54*, *TPD53* has often been used as a contrasting example. However, its persistent upregulation in multiple sample types across disease states and consistency in *in vivo* models suggest a much deeper and more important role for this overlooked gene which encourages more research in this direction(48–51).

Similarly, aquaporin 1 (*AQP1*) is an obligate water channel expressed in primary neurons, brain endothelial cells, and choroid plexus epithelial cells (involved in cerebrospinal fluid production) and is potentially involved in pain perception(52). Several studies have reported increased Aquaporin expression in the brain of AD patients, especially in regions adjacent to amyloid plaques, and it has been implicated in cerebral vasogenic edema observed in AD patients during anti-amyloid immunotherapy. Moreover, cortex astrocytes displayed increased expression of *AQP1* in low anti-amyloid immunoreactive regions around the senile plaques(53,54).

Therefore, upon observing these commonly altered genes in several diseases and sample types, we constructed Gene Ontology pathway enrichment and Gene-microRNA (miRNA) networks to investigate regulatory pathways and identify their regulatory components (miRNA) (**Figure 2)**. For example, our analysis further revealed that *TPD52L1* (*TPD53*) is mainly associated with positive regulation of stress-associated MAPK cascade and apoptotic process and *LMO4* with neural tube closure **(Figure 2A)** and identified four miRNAs interacting with the transcripts that can be targeted to alter the protein translation coded by these genes (**Figure 2B, C)** to test their regulatory effect in AD pathobiology.

**Figure 2:**
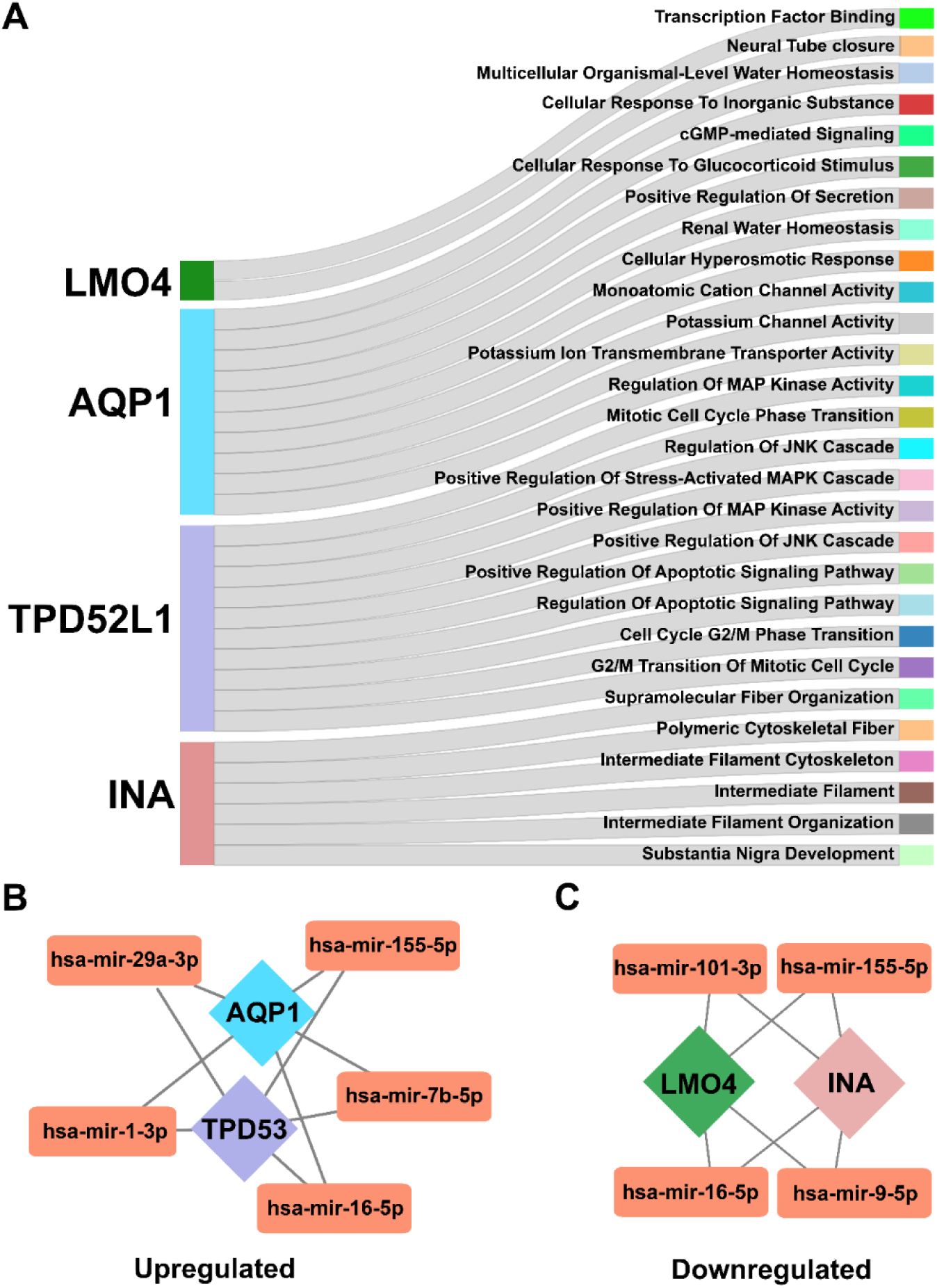
Pathways associated with genes that are commonly altered in different sample types across different diseases. (**A**) Sankey plot depicting the pathways enriched by the genes *AQP1*, *TPD52L1 (TPD53)*, and *INA*. (**B and C)** Gene–miRNA interaction networks depicting the miRNAs interacting with these genes.

iNSCs have been a common model for the *in vitro* modeling of neurodegenerative diseases, including AD(55). Our data also includes transcriptome data from iNSCs modeled for AD, DS, and MPS I. Comparison of the data across these diseases revealed three downregulated genes, *NETO2* (Neuropilin and Tolloid-like 2), *SFRP2* (Secreted Frizzled Related protein 2), and *CLSTN2* (Calsyntenin 2), and two upregulated genes *GDF15* (Growth Differentiation Factor 15) and *ADM2* (Adrenomedullin 2) in common (**Figures 1F, G**). Although we identified several genes commonly altered between AD and other diseases, we would like to highlight the importance of *NETO2* since it is also commonly downregulated across three AD sample types: brain samples, organoids, and iNSCs. The effect of *NETO2* on cognitive functions and its possible role in AD has never been studied, indicating that our analyses revealed a new gene associated with the genetic etiology of AD. Further research is needed to understand whether the decreased *NETO2* expression is beneficial (adaptive) or regulates the progression and pathogenesis of AD.

Overall, a comparison of transcriptional profiles of each disease across different sample types revealed commonly altered, previously unknown genes. Notably, downregulated *INA, NETO2*, and *TPD52L1* genes are commonly altered in both brain and organoid samples of AD, DS, and NPC suggesting them as potential molecular targets for examining their alteration in association with ameliorating the AD phenotype. Further research is needed to investigate the potential of these therapeutic strategies targeting the functions of these proteins on AD progression and pathogenesis. Although the identified target genes were low in number, we believe that these genes might represent the true molecular targets for AD and provide a foundation for future animal and clinical studies, leading to a better understanding of the molecular mechanisms of AD pathology and revealing diagnostic and therapeutic applications against common types of dementia.

### AD-like phenotype and accelerated brain aging in Niemann-Pick type C disease mouse model; a novel short-lived Alzheimer’s Disease model

NPC is a rare, prematurely fatal, inherited neurodegenerative disease that develops from lysosomal storage disorder and differs in several respects from Alzheimer’s disease (AD). However, intriguing parallels exist in the different pathological similarities of these two neurological disorders. For example, deposition of amyloid-β, hyperphosphorylated tau, neurofibrillary tangle formation, and neuronal cell death are associated with neurodegeneration. Additionally, abnormal cholesterol metabolism, cognitive decline, and severe dementia are pathogenic features of both NPC and AD(56–60). To this date, the underlying pathophysiological mechanisms are not yet well understood.

Unfortunately, as stated in the introduction, NPC is a rare disease, often leading to stillbirths or infantile deaths, meaning that the brain samples for this disease are severely limited, especially from adults and the elderly, as the lifespan reduction in juvenile-onset NPC is just as severe as infantile-onset NPC. In addition, there has not been a single report on transcriptomic data from the frontal cortex of NPC patient brains as the majority of work in the field is focused on the mitigation of the motor function loss that ultimately drives NPC-related mortality.

Considering these factors, we aimed to characterize the commonalities in disease phenotypes and molecular mechanisms involved in both NPC and AD pathogenesis. To achieve this, we leveraged the NPC1^mut^ mouse model (*Npc1*^tm(I1061T)Dso^), which is a crucial tool in the understanding of the pathobiology of NPC disease (14). This model, expressing the most common mutation in Npc1 protein seen in juvenile-onset NPC, faithfully recapitulates many aspects of human NPC disease. It presents a slower progression of neurodegeneration and a less severe disease phenotype, allowing for a longer longitudinal observation of disease progression. Importantly, it reliably demonstrates APP processing, dysregulation, and tau overexpression. However, it’s essential to acknowledge the potential drawback: motor coordination problems and ataxia can interfere with the model’s ability to participate in cognitive and behavioral tests. To address this, we conducted a minor feasibility study to characterize lifespan, onset of neurological impairment, and general reduction in motor function and coordination over time (**Figure 3**). As a primary observation, we identified a significant weight loss starting from week 9 (**Figure 3A**) and a sudden drop in percent survival between weeks 10 and 15 in NPC1^mut^ mice relative to wild-type mice. The median lifespan of the NPC1^mut^ mouse model was fifteen weeks (**Figure 3B**).

**Figure 3:**
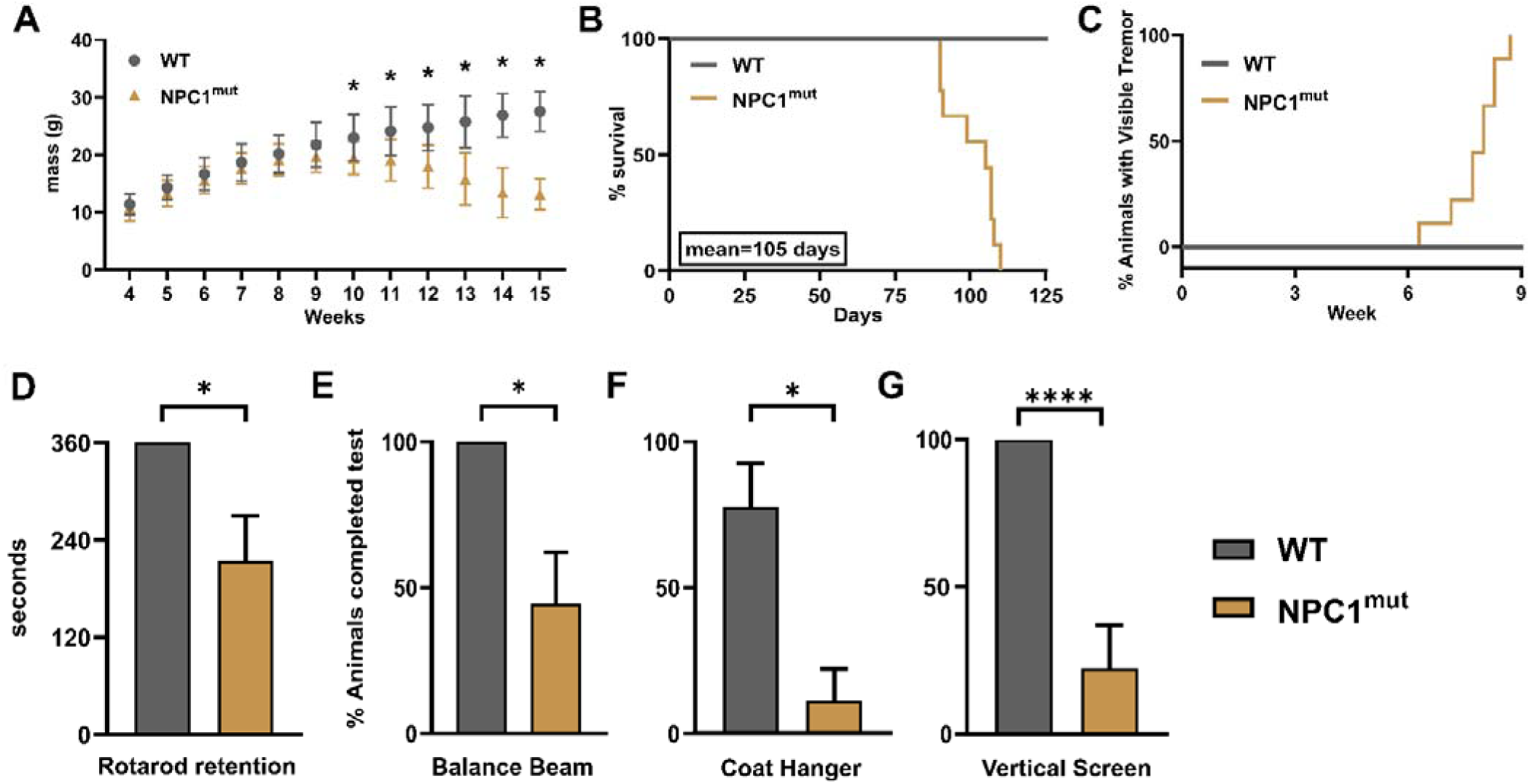
Minor Feasibility assessments to characterize motor function loss and ataxia in the NPC1^mut^ mouse model (both sexes) before the onset of neurological impairment. (**A**) Mass and (**B**) survival of WT and NPC1^mut^ mice. No WT animals died during the observation period of 20 weeks, and the median humane survival of NPC1^mut^ mice was 15 weeks (105 days). N=9 for all groups for the survival assay. N=22 for NPC1^mut^ mice and N=14 for WT mice for the weight loss study. Data in (**A**) represent mean + SD analyzed by mixed-effects model (REML) followed by Šídák’s multiple comparisons test to account for animals that died before week 15, *p≤0.05. (**C**) Mice were observed daily for the appearance of visible tremors. The median age at which NPC1^mut^ mice develop tremors was eight weeks. (**D – G**) Coordination testing of NPC1^mut^ mice at 13 weeks of age using (**D**) Rotarod testing at 15 RPM, and a battery of tests (**E – G**) developed specifically to monitor NPC1-related motor dysfunction as previously described. N=11 for all groups. (**D – G**) Data represents mean + SD and analyzed by t-test, * p≤0.05, **** p≤0.00005.

At a median age of eight weeks, NPC1^mut^ mice developed visible tremors and overt symptoms of ataxia despite retaining a mean retention time of over three minutes on the rotarod, even at thirteen weeks (**Figure 3C, D**). The entire cohort of NPC1^mut^ mice was able to balance themselves on the 2 cm round beam and traverse the beam of length 60 cm in less than three minutes, completing the test (**Figure 3E**). These results suggest that NPC1^mut^ mice retain requisite motor functions to assess cognitive abilities for at least five weeks after the first overt signs of neuropathy. We performed several tests specifically developed to determine *NPC1*-related motor dysfunction, as described in prior research(26). Surprisingly, however, these tests, which assess grip strength, climbing ability, and complex movements, suggest a significant neurological impairment in NPC1^mut^ mice (**Figure 3F, G**).

We observed slight sex-specific differences both in weight loss and age at which they develop visible tremors. Male NPC1^mut^ mice developed tremors faster than females by approximately a week. Similarly, they demonstrated initiation of weight loss two weeks before female mice, starting at 9 weeks, while female mice showed signs of weight loss starting from eleven weeks. These results demonstrate the underlying sex-specific differences in manifesting NPC1^mut^ phenotype, as evident from the different pathways enriched by DEGs from NPC1^mut^ versus WT female mice relative to NPC1^mut^ versus WT male mice (**Supplementary Figure 4).**

Next, to analyze the molecular commonalities between NPC and AD, we procured the NPC (*Npc1*^tm(I1061T)Dso^) mouse model, collected frontal cortex samples (n=3 for both sexes, total 6 samples for each background “WT versus NPC1^mut^” at 2 months old mice), and performed RNA sequencing (RNA-Seq) analysis (**Figure 4A, Supplementary File 2**). In comparison to the age- and sex-matched WT control, we found 714 downregulated for females and 75 downregulated genes for NPC1^mut^ male mice (*adjP* ≤ 0.05). We found that 39 downregulated genes were shared between male and female samples (**Figure 4B, Supplementary File 2**). There were 996 upregulated genes in female NPC1^mut^ mice and 117 upregulated genes in male NPC1^mut^ mice. Of these, 51 were common between both sexes. (**Figure 4C, Supplementary File 2**), further suggesting a sex-specific effect of *NPC1* mutation.

**Figure 4:**
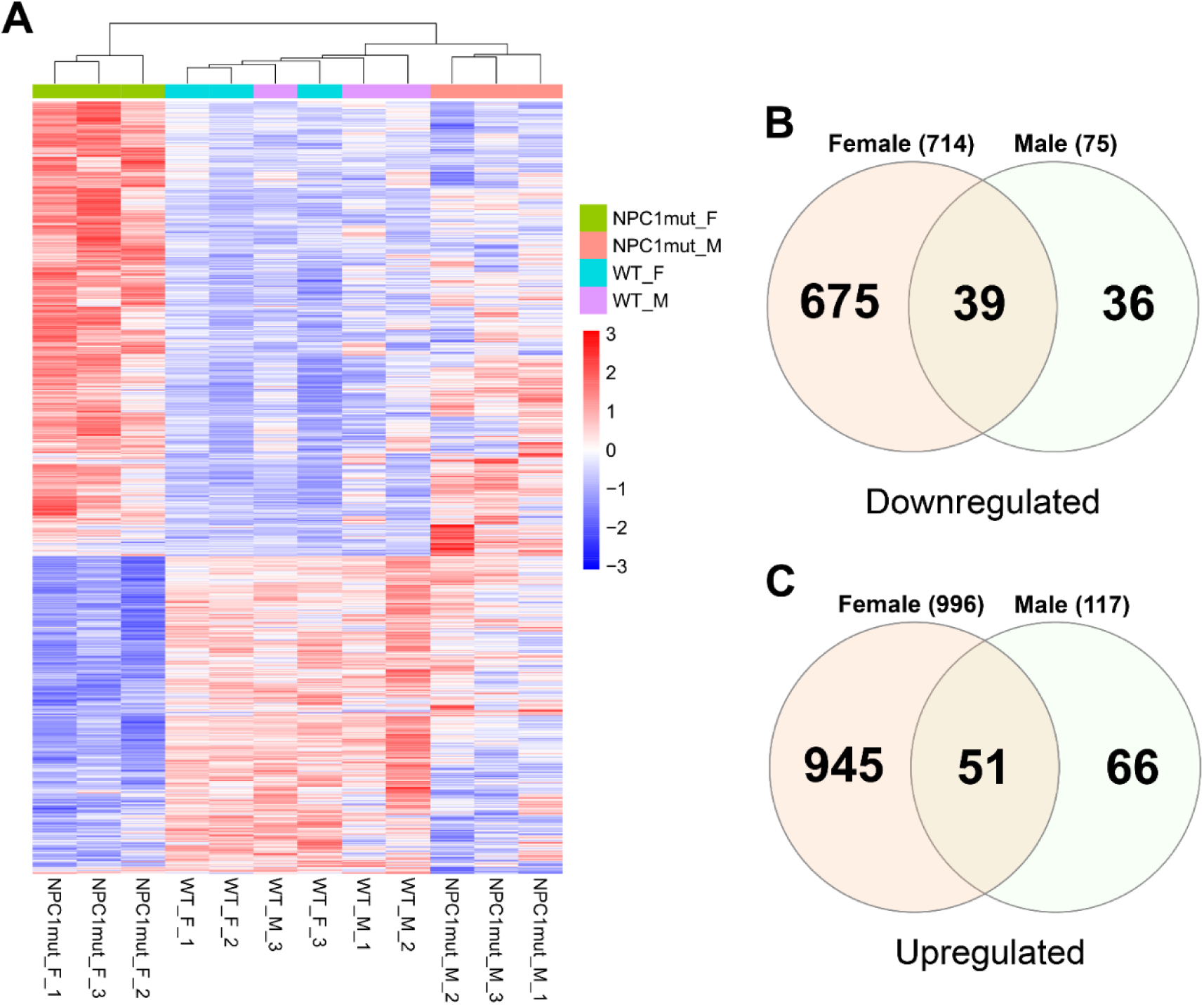
Sex-dependent gene expression changes in NPC1^mut^ mice. (**A**) The heatmap depicting the similarities and differences in differential expression of significantly altered genes in frontal cortex brain samples across NPC1^mut^ mice and age- and gender-matched WT controls. (**B and C**) Venn diagrams depict the commonality analysis results on differentially expressed genes from female and male NPC1^mut^ mice relative to age and gender-matched WT controls. (**B**) Among the 714 downregulated genes from female samples and 75 from the male mice, 39 genes are common. (**C**) Similarly, 51 genes are commonly upregulated between female and male mice samples. Supplementary File 2 contains the complete list of DEGs, normalized reads, p values, and the common gene list.

Then, we performed functional enrichment analyses for the DEGs explicitly identified for each sex. In female NPC1^mut^ mice, the downregulated DEGs significantly enriched in mRNA binding, processing, and splicing, while upregulated DEGs enriched in positive regulation of exon extension, astrocyte, and glial cell projection, glutamatergic synapse, glycosphingolipid biosynthesis, and focal adhesion (**Supplementary Figure 5A, B**). The data suggests that upregulated pathways are mostly compensatory mechanisms against neuronal damage, as discovered recently and associated with neuroinflammation(61–63). For the male NPC1^mut^ mice, the downregulated DEGs enriched in plasma membrane organization, acting binding, adherens junction, lactate dehydrogenase activity, and regulation of dendritic spine morphogenesis. The upregulated DEGs are enriched in amyloid-beta clearance by the cellular catabolic process, cell-cell contact zone, and axon terminus (**Supplementary Figure 6A, B**). Overall, this data shows sex-specific phenotypic effects of NPC disease in mouse models and portrays sex-specific significant differences in molecular changes in the frontal cortex.

Finally, to examine whether short-lived NPC1^mut^ mice are accompanied by accelerated brain aging and accumulation of mortality-associated molecular biomarkers, we utilized recently developed transcriptome-based clocks for the prediction of lifespan-adjusted age and expected lifespan(28). Aging is the progressive deterioration of cellular and organismal function, and age-dependent functional decline in cellular processes is linked to various neurodegenerative diseases, including AD(64–66). Our application of transcriptome aging clocks revealed accelerated brain aging and decreased expected lifespan (**Figure 5A, B**) in both male and female NPC1^mut^ mice. To further characterize cellular pathways that may drive the pro-aging phenotype of NPC1^mut^ mice, we performed gene set enrichment analysis (GSEA) of changes induced by *NPC1* mutation and established signatures of aging and mortality in mice and rats. We observed a significant positive correlation between functional changes associated with inhibited *NPC1* function, aging, and mortality (**Figure 5C**). In particular, we observed pro-aging upregulation of genes involved in interferon signaling, inflammatory response, MAPK signaling, and complement cascade in NPC1^mut^ mice, especially in females (**Supplementary Figure 7A, B**), along with pro-aging downregulation of genes associated with mitochondrial translation in both sexes (**Figure 5D**, **Supplementary File 3**). Our study represents a novel short-lived mouse model with accelerated brain aging, to serve as a reference for future application in fundamental and translational research of mammalian and more specifically human aging.

**Figure 5:**
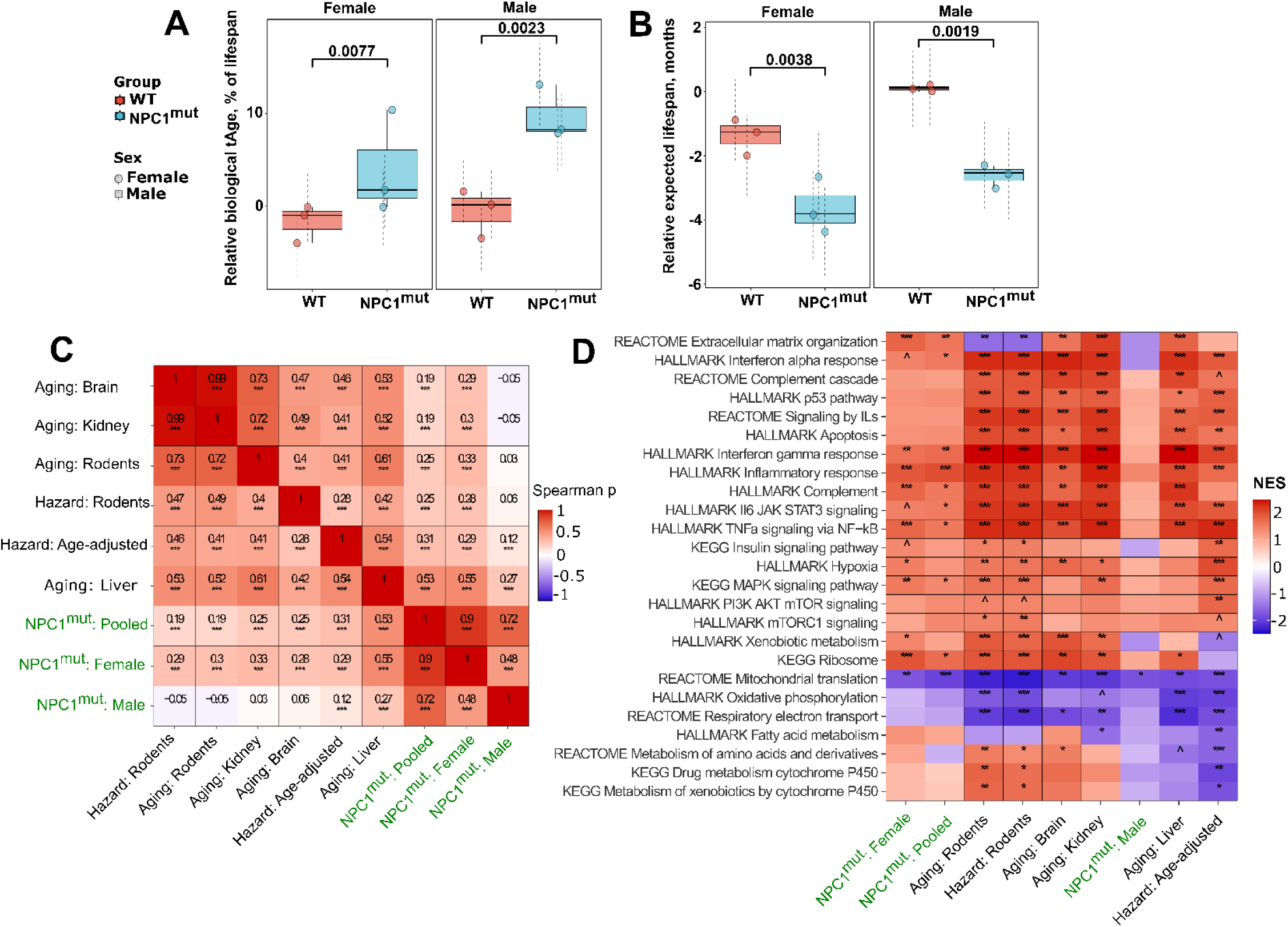
Association of gene expression changes in NPC1^mut^ mice with biomarkers of aging and mortality. Transcriptomic ages (tAges) and expected lifespans of control and NPC1^mut^ mouse brain samples were estimated with relative multi-tissue (**A**) lifespan-adjusted and (**B**) lifespan clock, respectively. Difference between groups was estimated for each sex. Data are mean ± standard deviation of posterior prediction. Labels reflect Benjamini-Hochberg (BH) adjusted p-values. (**C**) Spearman correlation between normalized enrichment scores (NES) of pathways associated with Npc1^mut^ and established signatures of aging and mortality. Signatures of aging and expected hazard are labeled in red, whereas signatures NPC1^mut^ mice are labeled in dark blue. (**D**) Functional enrichment (GSEA). Pathways enriched by gene expression changes induced in NPC1^mut^ mice (dark blue) from the current study as well as by established signatures of aging and mortality (red). Only functions significantly enriched by at least one signature (p adjusted < 0.1) are presented. The whole list of enriched functions is in Supplementary File 3. Benjamini-Hochberg p.adjusted < 0.1, *p.adjusted < 0.05, **p.adjusted < 0.01, ***p.adjusted < 0.001.

### Cross-comparison of molecular changes across NPC1^mut^ and AD mice

To characterize the efficacy of our NPC1^mut^ mouse as a short-lived AD mouse model, we performed commonality analysis on the DEGs of frontal cortex transcriptomic data obtained from the male and female APP/PS1 AD mouse model (NCBI GEO Accession number: GSE85162) at an appropriate time point (8 months) in which AD-like phenotype emerges. APP/PS1 mouse model is the most commonly utilized mouse model for AD research(15). In addition, multiple studies have shown that the *APP/PS1* model manifests the most robust gene expression signatures that significantly overlap with AD-associated co-expression signatures from human brains(46).

AD female mouse had 1,490 downregulated genes, and the AD male mouse model had 1,471 downregulated genes, of which 1,026 were common between female and male AD samples (**Supplementary Figure 8A, Supplementary File 4**). In addition, we identified 1,841 upregulated genes for females and 1,808 upregulated genes for male AD mice. Of these, 959 upregulated genes were common between male and female AD samples (**Supplementary Figure 8B, Supplementary File 4**). A comparison of sex-specific commonalities between AD and NPC1^mut^ mouse models revealed 57 commonly downregulated genes between NPC1^mut^ and AD females (**Supplementary Figure 9A, B and Supplementary File 5**) and only 9 genes commonly downregulated between NPC1^mut^ and AD male samples (**Supplementary Figure 9A, C and Supplementary File 5**). A cross-comparison of DEGs found 62 upregulated genes in female samples (**Supplementary Figure 10A, B and Supplementary File 5)** and 7 upregulated genes in male samples were common between AD and NPC1^mut^ mouse samples (**Supplementary Figure 10A, C and Supplementary File 5**).

The enrichment analysis for these common genes has uncovered novel pathways. The downregulated common genes were associated with the spliceosome and actin filament binding processes, which are crucial for maintaining the integrity of neuronal structure (**Supplementary Figure 11A**). In contrast, the upregulated genes are primarily linked to neurogenesis, neuronal differentiation, and synaptic response (**Supplementary Figure 11B**). This seemingly paradoxical finding is in line with recent research that has observed the upregulation of synaptic pathways in early AD(67). Moreover, the upregulated neurogenesis and neuronal differentiation appear to be compensatory mechanisms triggered by neuronal loss from AD progression, providing a fresh perspective on our results(61,68). It is important to note that the low number of common genes identified for the male samples of AD and NPC1^mut^ limited the enrichment analyses from yielding any significant terms.

Overall, these data showed that compared to the NPC1^mut^ mice, AD mice had a higher number of transcriptional changes in both male and female brain frontal cortex samples, and a significant number of genes were commonly altered in both models, further suggesting some of the identified genes and pathways might commonly be associated with disease pathobiology.

### Cross-comparison of human brain samples revealed significant disease similarities between AD and NPC at the transcriptional level

To further characterize whether the NPC1^mut^ mouse model can better recapitulate the molecular changes of human disease conditions, we performed a similar commonality analysis by comparing the transcriptome data from human AD brains to AD and NPC1^mut^ mouse frontal cortex brain samples separately. In our comparison, we searched whether any significantly altered human genes were detected in the combined (female and male) DEG list of AD or NPC1^mut^ mice samples (**Figure 6**). Our analyses revealed that 1.71% of the downregulated (57 orthologous genes out of 3,330 [1,490 genes from female and 1,840 genes from male]) (**Supplementary Figure 12A, B**) and 1.01% of the upregulated genes (44 orthologous genes out of 4,377 [1,759 genes from female and 2618 genes from male]) from AD mice samples were commonly detected in human AD brain samples (**Supplementary Figure 12C, D and Supplementary File 6)**. Among them, *RSPO3, TPD52L1*, *and ZNF248* were commonly upregulated. The other four genes, *HLA-A, SCG3, THY1, and IGFBP2*, were commonly downregulated between human AD organoids and human orthologs of mouse DEGs. Of these, the importance of the genes *TPD52L1 and HLA-A* was discussed elsewhere, and their differential expression in AD mouse samples akin to human samples strengthens the necessity for further research into these genes and their encoded proteins.

**Figure 6:**
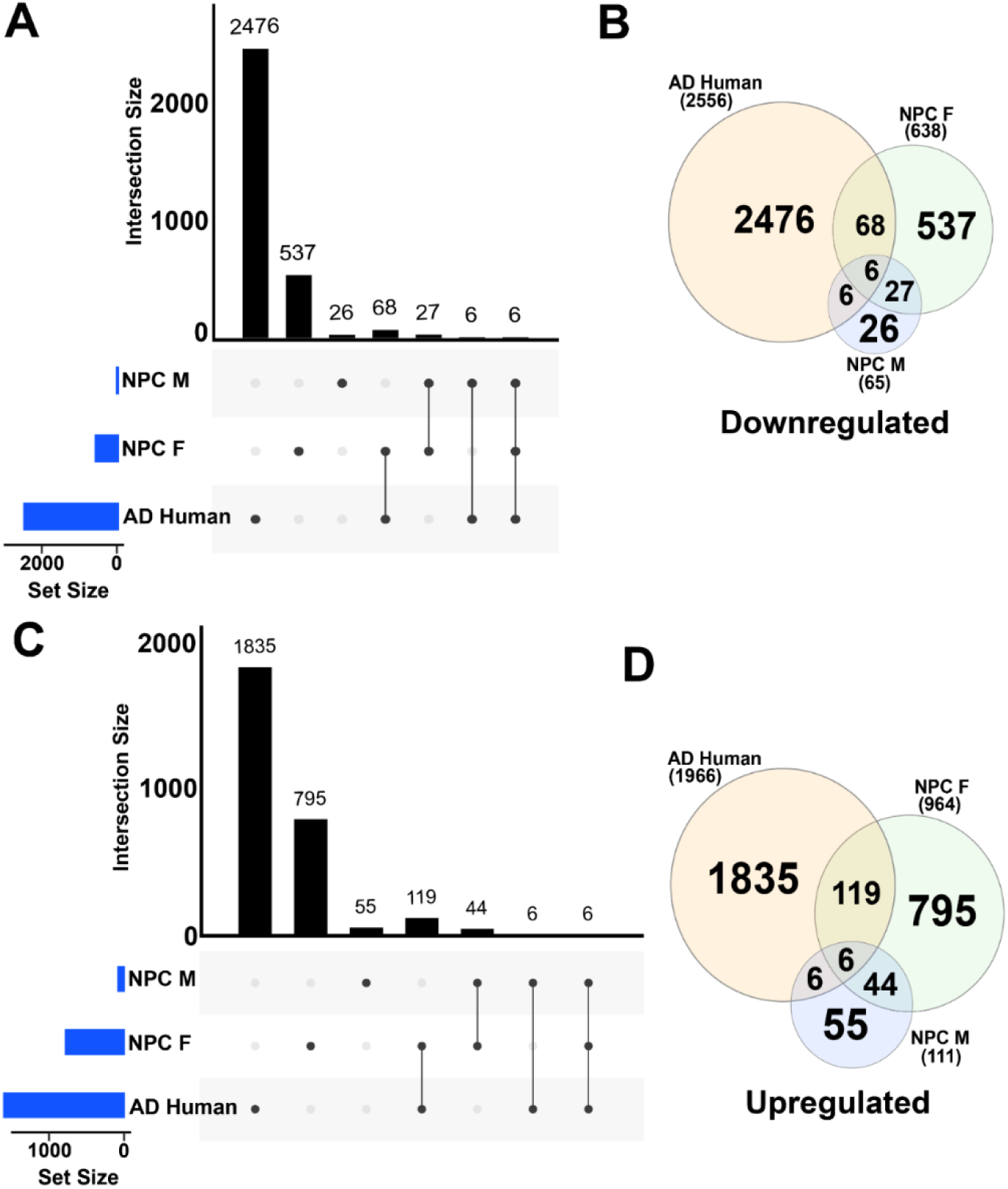
Cross-comparison of gene expression commonalities between frontal cortex samples of NPC1^mut^ mice and human AD brain samples. Upset plot (**A**) and (**B**) Venn diagram depicting the downregulated genes common to AD brain samples from humans and human orthologs from the NPC1^mut^ mouse model. Only 6 genes; *COL5A1*, *DLK2*, *ANKRD4*, *MATK*, *MIAT*, and *MICAL2*, are downregulated in all sample types. In total, 68 downregulated genes are common to AD brain samples and NPC1^mut^ female mouse models. In contrast, only 6 downregulated genes were common to the AD brain and NPC1^mut^ male mouse samples. Upset plot (**C**) and (**D**) Venn diagram depicting the upregulated genes common to AD brain samples from humans and human orthologs from the NPC1^mut^ mouse model. There are 6 upregulated genes, *PCDHGB3*, *ABCA1*, *CHD7*, *FLT1*, *RNF213*, and *MYO10*, common to all sample types. Similarly, six upregulated genes are common in AD brain samples and NPC1^mut^ mouse male samples. In total, 119 upregulated genes are common in human AD brain and NPC1^mut^ mouse female samples. The complete list of the commonly altered genes can be found in Supplementary File 6.

On the other hand, 10% of the downregulated (80 orthologous genes out of 792 [717 genes from female, 75 genes from male]) (**Figure 6A, B)** and 11.5% of the upregulated genes (131 orthologous genes out of 1126 [1007 genes from female and 119 genes from male]) from NPC1^mut^ mice samples were commonly detected in human AD brain samples (**Figure 6C, D and Supplementary File 6)**. Examining the commonalities revealed that *MIAT, MICAL2*, *COL5A1*, *DLK2*, *ANKRD24*, and *MATK* commonly downregulated in all three samples: AD brain samples, NPC1^mut^ female mice samples, and NPC1^mut^ male mice samples. Another six genes, *PCDHGB3*, *ABCA1*, *CHD7*, *FLT1*, *RNF213*, and *MYO10,* were commonly upregulated in all three samples. It should be noted here that many of these identified genes have never been investigated in the context of AD progression and pathology. For example, the *MIAT* gene (myocardial infarction-associated transcript), commonly downregulated in human AD brain samples and female and male NPC1^mut^ mice samples, encodes a long non-coding RNA, conserved across mice and humans that promotes neurovascular remodeling in the eyes and the brain(69) whose pathways have been associated with myocardial infarction, schizophrenia, age-related cataract, ischemic stroke, and cancers in humans and retinal cell fate determination in mice(70). This gene and its function have also been implicated in regulating the formation of advanced atherosclerotic lesions and destabilizing the plaques(71). Recent research highlights the ability of exosome-derived *MIAT* to improve cognitive dysfunction in vascular dementia rat models, suggesting the adverse effects of its downregulation, thereby proving the importance of our results(72). This gene is downregulated in AD mice and human AD organoid samples, indicating an important regulatory function in AD pathology.

We performed pathway enrichment analysis on the genes common across AD human and NPC1^mut^ female and male mice frontal cortex brain samples. We found several important pathways, like MAPK and PI3K – Akt signaling pathways, Apolipoprotein binding, and regulation of actin cytoskeleton, the importance of which was discussed previously. We also generated protein-protein interaction networks for the genes involved in the enrichment of some of the important pathways from the aforementioned pathway enrichment to highlight the interactions among those genes. (**Figure 7A, B and Supplementary File 7)**

**Figure 7:**
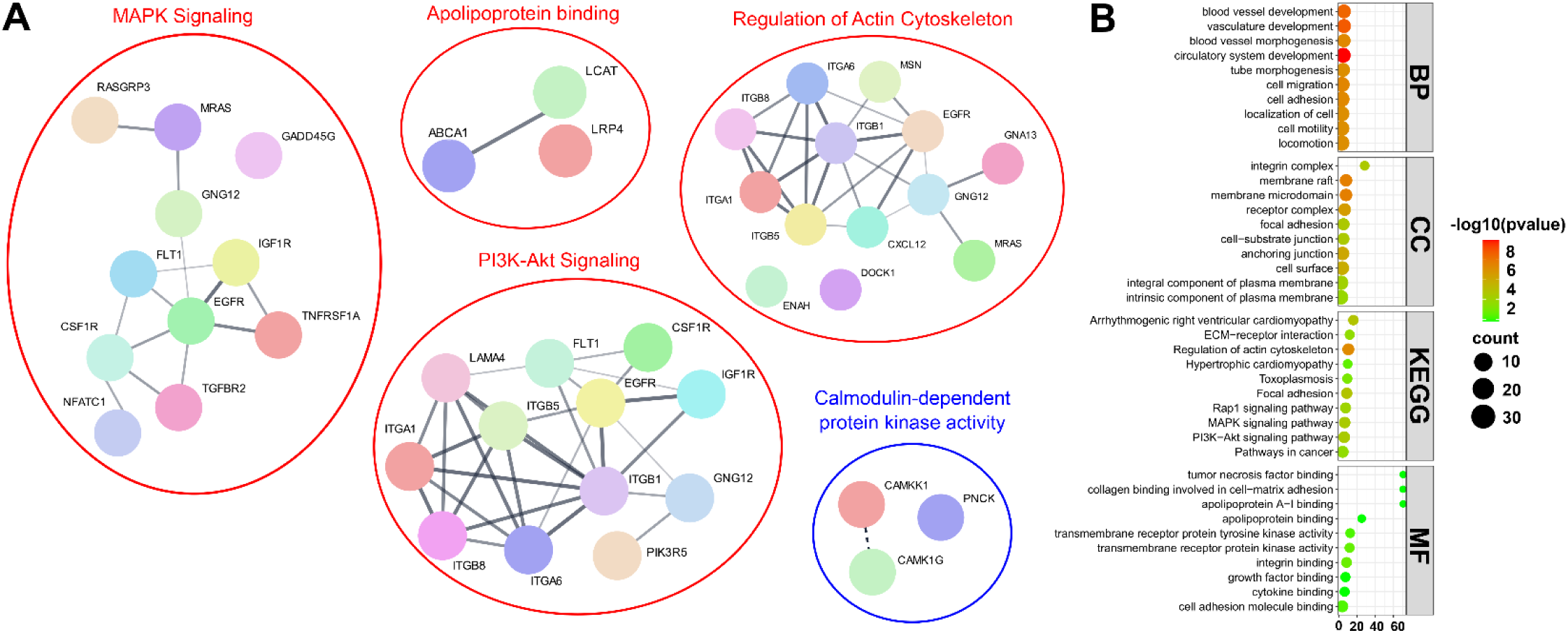
Protein-protein interaction networks and pathway enrichment analysis for the common differentially expressed genes between AD human brain and NPC1^mut^ female and male mice brain samples. (**A**) Protein-protein interaction network and enriched pathways for the commonly up (Red) and downregulated (Blue) genes from Figure 8. MAPK signaling, PI3K – Akt signaling, Apolipoprotein binding, and Regulation of actin signaling pathways are among the enriched pathways for the upregulated genes. The thickness of the connecting lines indicates the confidence of the interactions between the connected proteins. The enrichment analyses revealed a single significantly enriched pathway associated with Calmodulin-dependent protein kinase activity. (**B**) Significantly enriched pathways were identified with GO and KEGG pathway terms for commonly upregulated genes in AD brain samples and the human orthologs of upregulated genes from male and female NPC1^mut^ mice brain samples. Certain pathways, such as actin cytoskeleton regulation, apolipoprotein binding, MAPK signaling, PI3K – Akt signaling, and Toxoplasmosis, are commonly enriched across different categories, reinforcing the importance of these pathways.

This data strikingly suggests that even though frontal cortex samples from NPC1^mut^ mice have fewer altered genes (compared to the APP/PS1), the higher number of overlapped genes compared to the human AD brain samples, especially considering the lower overlap of other human cell-derived sample types such as iNSCs or organoids, indicates that NPC1^mut^ mice better recapitulate AD. This fact is further reinforced by a meta-study published in 2020 by Wan et al. which compared AD human brain transcriptome with AD mouse models and found their ability to represent human AD lacking due to various reasons: most of them are based on single penetrant gene mutations from rare familial autosomal dominant forms of the disease, they are unable to present clinically-relevant pathologies seen in humans, and while human brain autopsies show multiple mixed pathologies mice models only exhibit a single homogenous pathology associated with its respective mutation (46).

## Discussion

Alzheimer’s disease is a complex neurodegenerative disorder with multiple entangled causes, many of which were only recently discovered or not studied enough. There are many complicated reasons for manifesting AD phenotype, ranging from NFTs and amyloid plaques to hypoxia and glucose metabolism impairment. With hypoxia (73), lysosomal storage disorders (74), and impaired glucose metabolism (75) serving as the new markers of AD symptoms, research on AD neurodegeneration must shift to a non-conventional direction instead of solely focusing on amyloid accumulation. Hence, we put forth this study as an attempt to steer AD research toward congenital disorders with an inherent risk of neurodegeneration and help bring new perspectives by analyzing molecular commonalities among them. In our quest, we found several genes, transcription factors, and miRNA interactors to construct an unknown regulator network of AD. We believe our study is a forerunner in understanding the underlying mechanism for increased AD risk in multiple congenital disorders to identify genetic targets for pharmacological applications to ameliorate AD phenotype. As such, our study can lay the foundation for the inclusion of congenital disorders related to dementia and neurodegeneration in AD research as well.

Among the genes we identified, *LMO4*, a transcription regulator and known differentiation repressor, was downregulated in AD brains, potentially to facilitate compensatory neuronal differentiation to ameliorate neuronal damage resulting from Neurofibrillary tangles (NFTs) and Amyloid plaques (40). *LMO4* also regulates calcium release and synaptic plasticity, core neural functions affected by AD (76). Hence, its downregulation in our study is understandable. However, observing the same reduction in NPC organoid samples indicates that its downregulation is not a simple coping mechanism manifesting after AD but also an early indicator of the risk of AD, elevating its importance. Similarly, transcriptional upregulation of *AQP1* across different diseases suggests an association between *AQP1* function and AD pathology. Its role in brain edema and traumatic brain injury is well documented (52,77). *AQP1* is also implicated in many neurological disorders, including AD (53). It is mainly thought to be expressed during or after the manifestation of disease phenotype, but its upregulation in the early stages of MPS I says otherwise. It is highly likely that its expression coincides with the emergence of the AD phenotype and exacerbates its pathogenesis, so regulating its expression may help reduce brain damage in patients. Organoid data also showed the importance of genes such as *QKI* and *RHOB*, associated with myelinization (78), RNA binding, and DNA damage-induced apoptosis (79), all crucial processes required to curb neurodegeneration, which are downregulated in both DS and NPC. The mammalian gene *QKI* (Quaking) encodes an evolutionarily conserved RNA-binding protein that post-transcriptionally regulates myelinization and developmental genes in the brain (80). It is expressed in glial cells and implicated in several neurological disorders, including AD. Based on recent evidence, glial cells play a crucial role in sporadic AD. *QKI* is necessary for oligodendrocyte development and myelinization (80). It is expressed in glial cells, which play a crucial role in sporadic AD, and implicated in several neurological disorders. The protein encoded by this gene is highly expressed in the murine brain, mainly in astrocytes, suggesting its importance in the central nervous system as well (81). It was downregulated in AD, as expected, but also in DS and NPC suggesting its downregulation may not be a consequence but one of the causes of AD pathogenesis. The downregulation of this gene in the APP/PS1 AD mouse model was observed as expected, further bolstering the importance of studying this underrepresented gene in AD.

*INA* encodes α-Internexin, an intermediate neuronal filament (41) implicated in neurodegenerative disorders (82) and recently discovered to be a crucial factor in Neuronal Intermediate Filament Inclusion Disease (NIFID) (43). It is found prominently in NIFID inclusions but a relatively minor product in inclusions or plaques of other neurodegenerative disorders (44). It is one of the first filament proteins expressed in neurons of the developing nervous system after mitosis. In adults, its expression is limited to mature neurons. Despite the knowledge of its downregulation in AD, there have not been many studies on *INA* or other neuronal filaments. Hence, its pronounced downregulation in DS and NPC (the diseases with the highest correlation to and risk for AD) in addition to AD is a crucial result. Its downregulation in NPC suggests a potential causative role for AD phenotype. Rho-related GTP-binding protein RhoB, encoded by *RHOB*, is a mediator of apoptosis following DNA damage and is involved in the intercellular trafficking of many proteins (83). One of its primary interactors in neurons is the microtubule-associated Associated Protein Tau (*MAPT*), a neurofibrillary tangle protein (78). RhoB is associated with aging, neurodegeneration, and traumatic brain injuries. Unlike AD, where there is a marked decrease in RhoB (84), RhoA and RhoB levels rise in traumatic brain injuries (85). As evident from the study on proapoptotic properties of RhoB in knockout mice upon doxycycline-mediated DNA damage by Kamasani et al., many of RhoB targets are genes associated with aging, oxidative stress in the brain, and AD, including amyloid precursor protein (*APP*), *CAV1*, and *SST* (one of the downregulated genes from frontal cortex samples) (79). They proposed that RhoB-mediated apoptosis may influence AD, as many of its targets include proteins like Clic1 (a chloride ion channel), and genes like *NAP1*, and *ROCK* associated with *APP* processing or amyloid-mediated toxicity (78). Hence, its downregulation in NPC and DS organoids suggests the importance of this gene in the initiation and progression of AD phenotype. It also strengthens the necessity of further research on this gene, its targets, and its interactions. Finally, *TPD52L1* (Tumor protein D52 Like 1) is commonly upregulated across different sample types of AD and in brain samples and organoids across AD, DS, and NPC diseases. Its upregulation in AD mouse samples suggests a potential causative role for AD pathobiology.

Despite their seeming lack of correlation, different sample types highlighted diverse aspects of AD risk in various congenital diseases, and the genes common across two or more of them emerged as biomarkers for predicting AD risk before manifestation. This lack of correlation across sample types shows the limitations of different sample types but also gives a glimpse into the complexity of AD and the necessity for incorporating more research into congenital diseases associated with neurodegeneration. To conclude, we identified the genes *AQP1, TPD52L1, QKI, RHOB, PTPRO, ASNS, DDIT3, TNMD, NETO2, ZNF248, PCDH1, STON2, SHISA2, SPON2, GDF15, INA*, and *LMO4*, miRNA interacting with the genes, namely hsa-mir-1-3p, hsa-mir-16-5p, hsa-mir-101-3p, hsa-mir-9-5p and hsa-mir-155-5p by comparing AD transcriptomic data with transcriptomic data from various samples of congenital disorders with increased AD risk, namely, NPC, DS, MPS I (**Supplementary Table 4**). While some of these genes have been associated with AD before, most were never implicated in neurodegeneration. We believe a consistent characterization of these genes across different samples and disease types highlights their importance for understanding AD biology and identifying them as a target for pharmaceutical approaches.

Perhaps one of the most significant contributions of our study is characterizing the *Npc1*^tm(I1061T)Dso^ mouse model as a short-lived model with an accelerated brain aging phenotype. Although short-lived mouse models with accelerated aging have already been characterized and commonly used for aging research, such as Hutchinson–Gilford progeria syndrome (HGPS), studies showed that this mouse model shows negligible changes in gene expression and does not reveal significant pathology to the AD brain (86). Our comparative transcriptomics and behavioral assays indicate NPC1^mut^ mice can be used to investigate the mechanisms underlying normal and pathological brain aging. Strikingly, in comparison to the similarities between the APP/PS1 mouse model and AD human brain samples, the number of commonly altered genes between the frontal cortex of the NPC1^mut^ mouse model and human AD brain samples was ∼10 times higher. In addition, commonalities among significantly enriched pathways suggest that NPC1^mut^ mice can serve as a potential short-lived *in vivo* model for AD research and for understanding molecular factors affecting brain aging. In fact, our aging signature analyses identified inflammation and mitochondrial dysfunction as major drivers of accelerated brain aging, further validating that NPC1^mut^ mice physiologically recapitulate aging hallmarks.

Among the genes commonly altered in AD human brain and NPC1^mut^ mouse brain were *MIAT, MICAL2*, *COL5A1*, *DLK2*, *ANKRD24*, and *MATK* commonly downregulated in all three samples: AD brain samples, NPC1^mut^ female and male mice samples. Another six genes, *PCDHGB3*, *ABCA1*, *CHD7*, *FLT1*, *RNF213*, and *MYO10*, were commonly upregulated in AD brain samples, NPC1^mut^ female and male mice samples. A surprising observation from this analysis was the upregulation of the gene *NPC2* in both AD brain samples and NPC1^mut^ female mice brain samples. While it is understandable in NPC1^mut^ mice as compensation against dysfunctional *NPC1*, observing the increase in *NPC2* expression in human AD brain samples reinforces the need for studying AD in the context of congenital disorders like NPC. Furthermore, the decreased expression of *MICAL2,* which catalyzes the selective oxidation of Met 44 and Met 47 residues, thus facilitating F-actin depolymerization (87), was shown in the synapse in adolescent APP/PS1 mice compared to WT (88). Considering that Aβ mediated F-actin depolymerization and loss of dendritic spines are associated with memory deficits in the early stage of AD, decreased *MICAL2* expression might be adaptive to compensate for F-actin loss in the AD brain.

In summary, we can conclude that the disease similarity approach by cross-comparing different biological data across diverse sample types and diseases was able to reveal unknown regulators and potentially identify new therapeutic targets to alleviate AD pathology. In addition, despite the large amount of research performed in animal models of AD, our results raise the exciting possibility that the NPC1^mut^ mouse model can serve to rapidly test the efficacy of interventions/combination of specific interventions to counter aging and AD. A systematic analysis focusing on the biological function and interactions of identified genes can provide valuable insights into understanding molecular features in AD and provide potential targets for developing pharmacological interventions. We believe that our characterization of the short-lived NPC1^mut^ mouse model as a robust model for aging and AD studies will further accelerate the research in these fields and offer a unique model for understanding the molecular mechanisms of AD from a perspective of accelerated brain aging.

## Author Contributions

Conceptualization; Kaya A., and Newton J., Methodology and Data Curation; Kaya, A. Newton J., Tyshkovskiy A., Oz N., Abyedah M., Gujjala VA., Formal Analysis; Tyshkovskiy A., Abyedah M., Gujjala VA., Castro JP., Newton J., Writing – Original Draft; Kaya A., and Newton J., Gladyshev VN., Castro JP., Gujjala VA. Animal Experimentation and Tissue Acquisition; Newton J., Kaya A., and Klimek I.

## Acknowledgments

We would like to acknowledge the generous funding provided by the NIA/NIH (1K01AG060040). Studies performed by J. N. were funded by the NICHD/NIH (5R00HD096117). The funders had no role in study design, data collection, and interpretation, or the decision to submit the work for publication. Services in support of the research project were provided by the VCU Massey Comprehensive Cancer Center Transgenic/Knockout Mouse Shared Resource, supported, in part, with funding from NIH-NCI Cancer Center Support Grant P30 CA016059.

## Data availability

All data generated or analyzed during this study are included in the manuscript and supporting files. RNA-seq data from the frontal cortex samples of *Npc1*^tm(I1061T)Dso^ mouse is deposited to the NCBI Gene Expression Omnibus (GEO) with accession number GSE266485.

---

**Figure S1:**
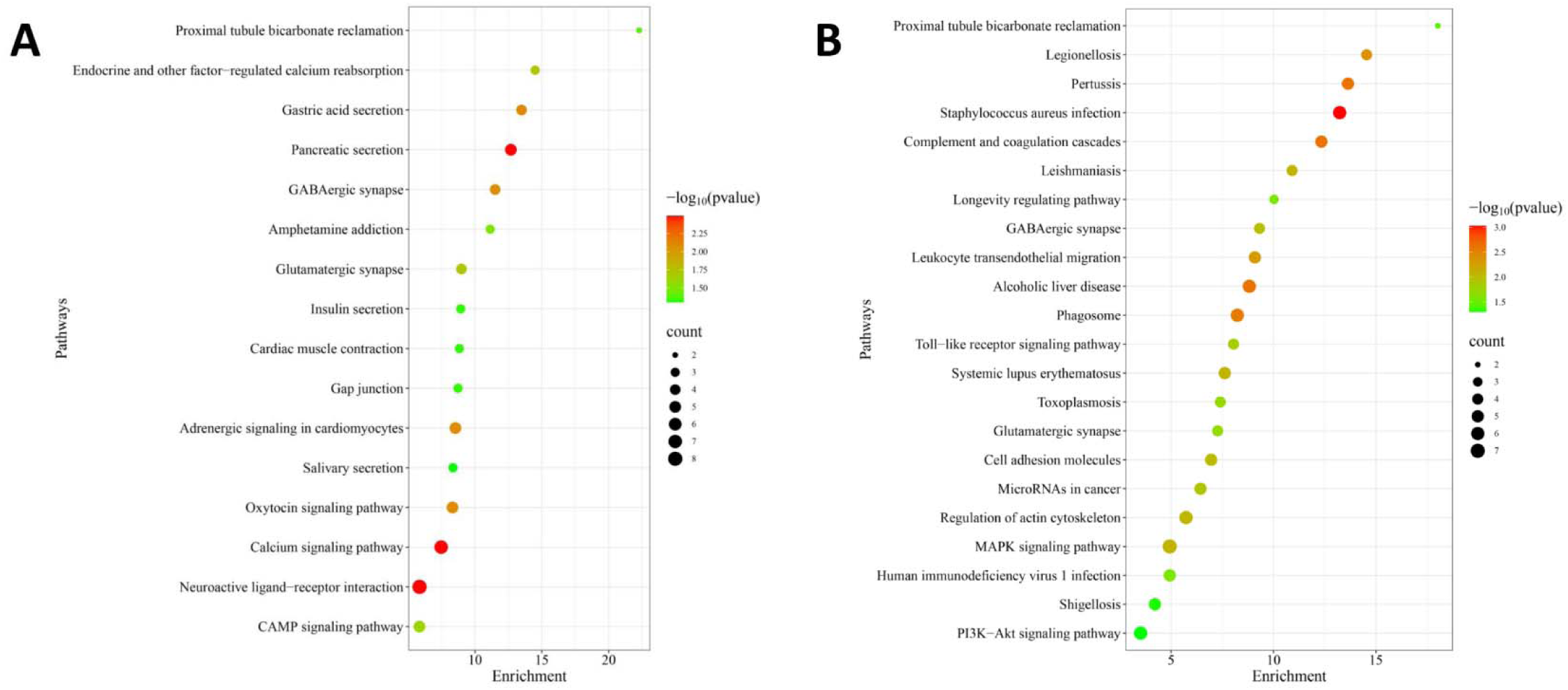
GO enrichment terms for commonly altered genes between AD and DS brain samples. Enrichment bubble plots depicting the pathways enriched by the differentially expressed genes shared between frontal cortex samples of human AD and DS brain samples. (**A**) Pathways enriched for the 96-downregulated and (**B**) for the 117-upregulated genes are shown.

**Figure S2:**
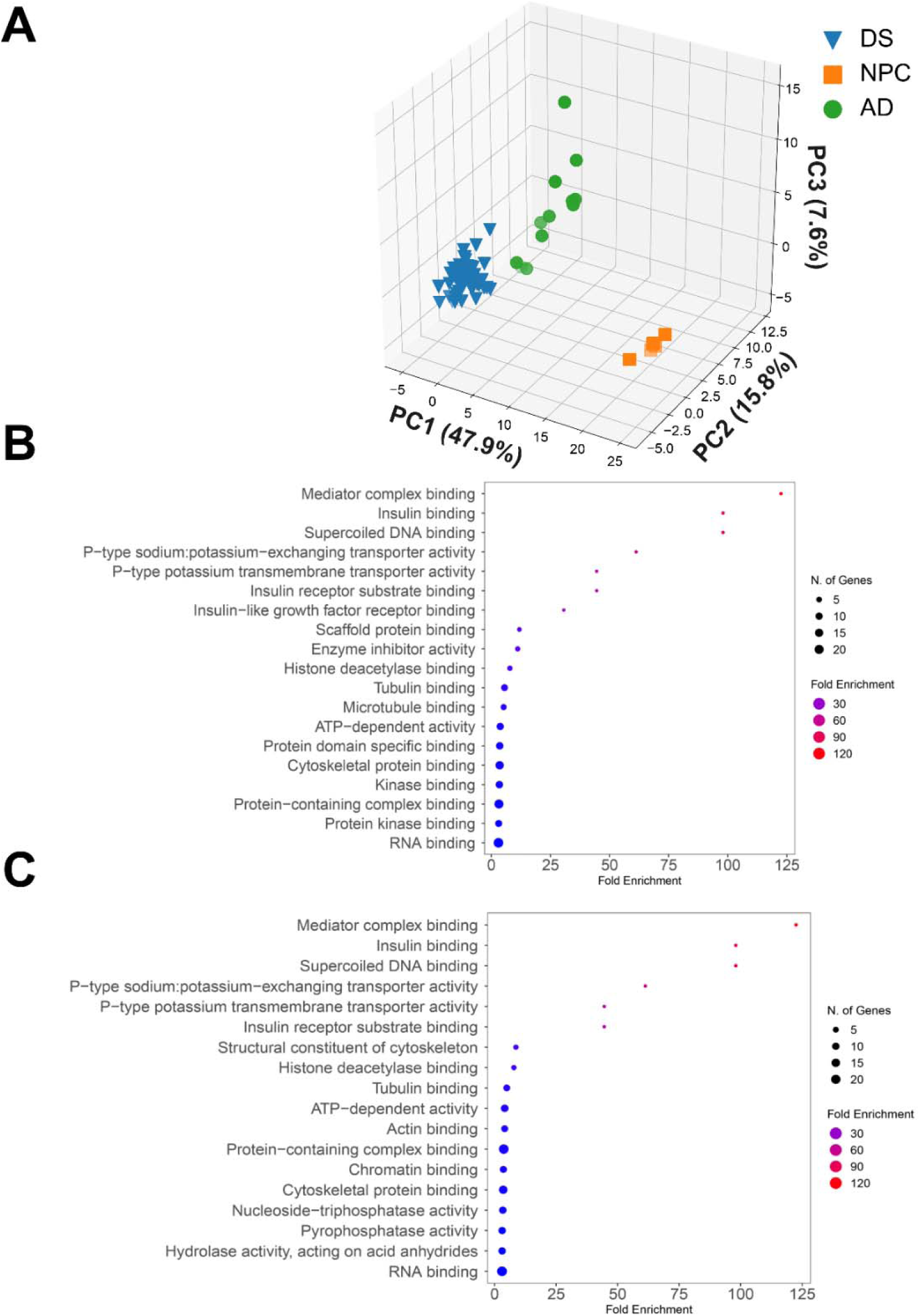
Sample distribution and differentially regulated pathways across NPC, AD, and DS organoid samples. (**A)** The PCA scatter plot depicting the sample variability between NPC (orange), AD (green), and DS (blue) based on gene expression signatures. PC1 shows the significant similarities between AD and DS samples and their distinction from NPC samples. PC2 shows less variability among samples of all three disease types. (**B and C)** The GO Enrichment (Molecular Function) terms are depicted with the dot plot for the pathways enriched for the top 50 positive and negative genes that explain the maximum variance for the PC1 and PC2, respectively. The complete list of genes used for enrichments based on the PCA scores can be found in Supplementary File 1.

**Figure S3:**
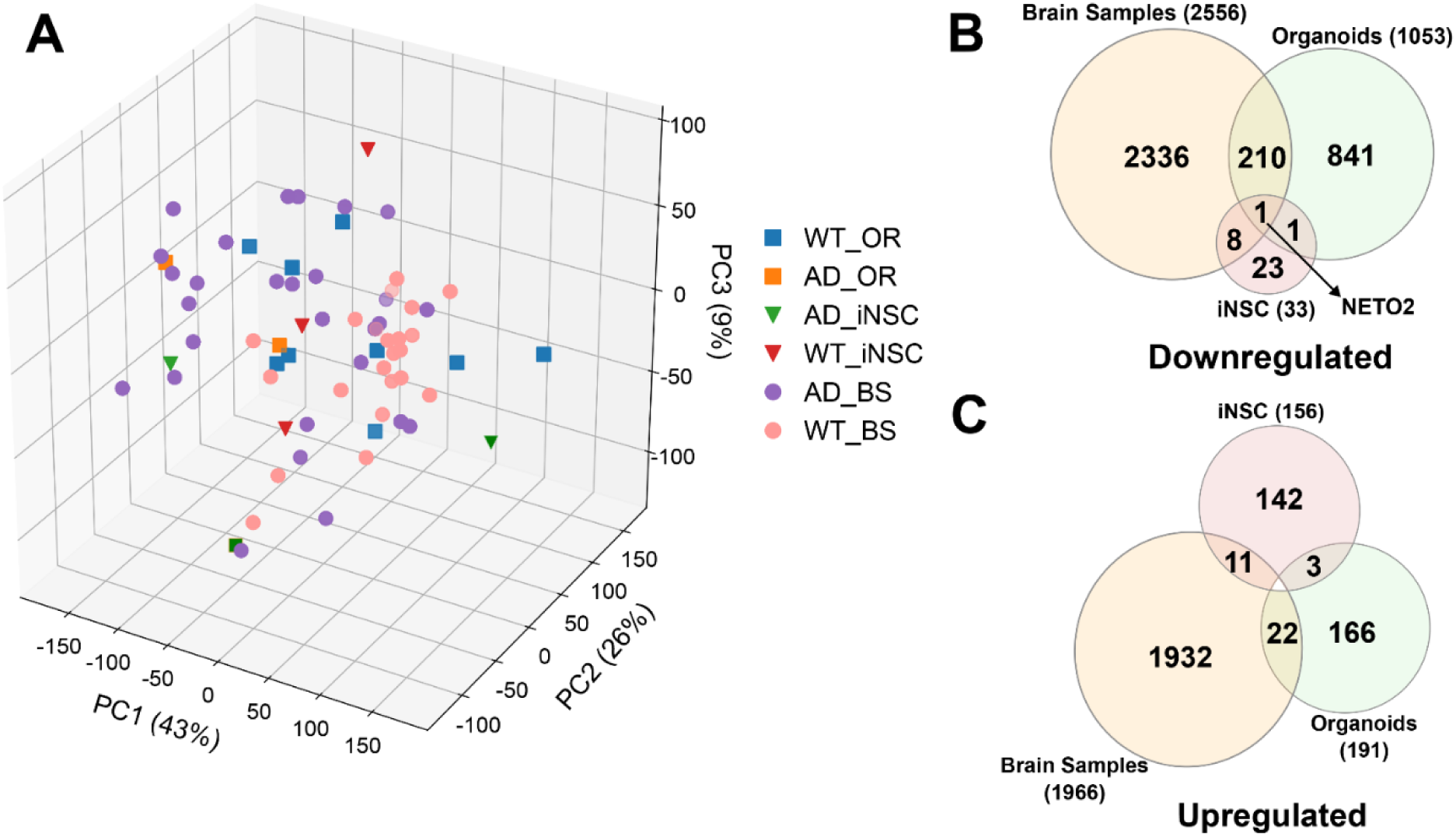
Cross-comparison of gene expression changes across different models of AD. **(A)** Principal Component Analysis (PCA) on transcriptomic data for AD from various sample types against the respective wild-type controls. (**B and C)** Venn diagrams representing the common DEGs across different sample types for AD. One gene, *NETO2*, is commonly downregulated across three sample types. There are no genes commonly upregulated across all sample types. Based on the number of common DEGs, the organoid model has shown the highest correlation with brain samples. This data concludes that none of the *in vitro* sample types can fully recapitulate the molecular signatures of Alzheimer’s brain at the transcript level.

**Figure S4:**
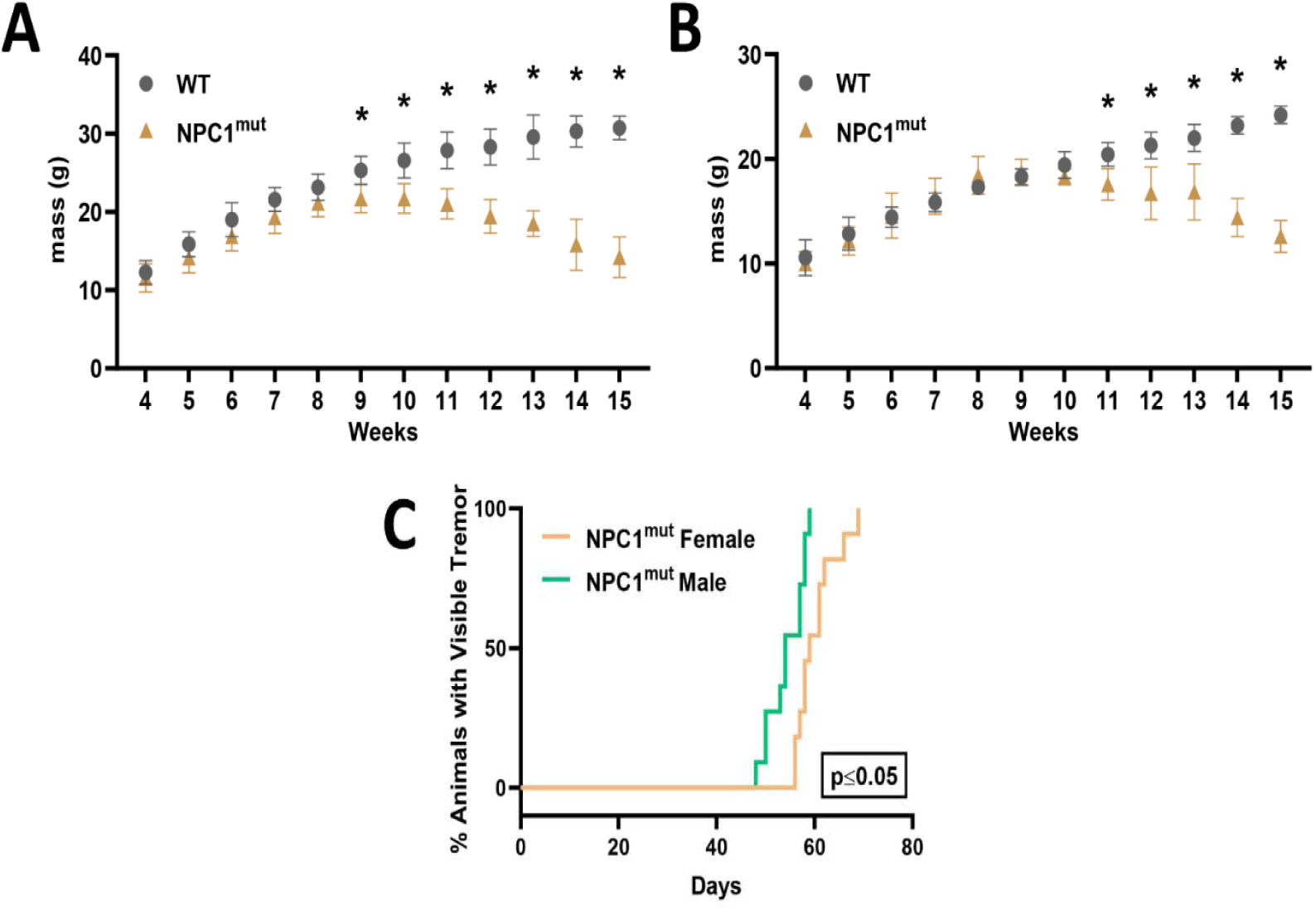
Minor Feasibility assays to characterize the motor function loss and ataxia in NPC1^mut^ mouse model (sex-specific) before the onset of neurological impairment. (**A**) Male and (**B**) Female weight-loss assessment of WT and NPC1^mut^ mice. N=11 for NPC1^mut^ mice and N=7 for WT mice. Data in (**A** and **B**) represent mean + SD analyzed by mixed-effects model (REML) followed by Šídák’s multiple comparisons test to account for animals that died before week 15, *p≤0.05. (**C**) Mice were observed daily for the appearance of visible tremors. The median age at which male NPC1^mut^ mice develop tremors was seven weeks (48 days). In contrast, the median age at which female NPC1^mut^ mice develop tremors was eight weeks (58 days).

**Figure S5:**
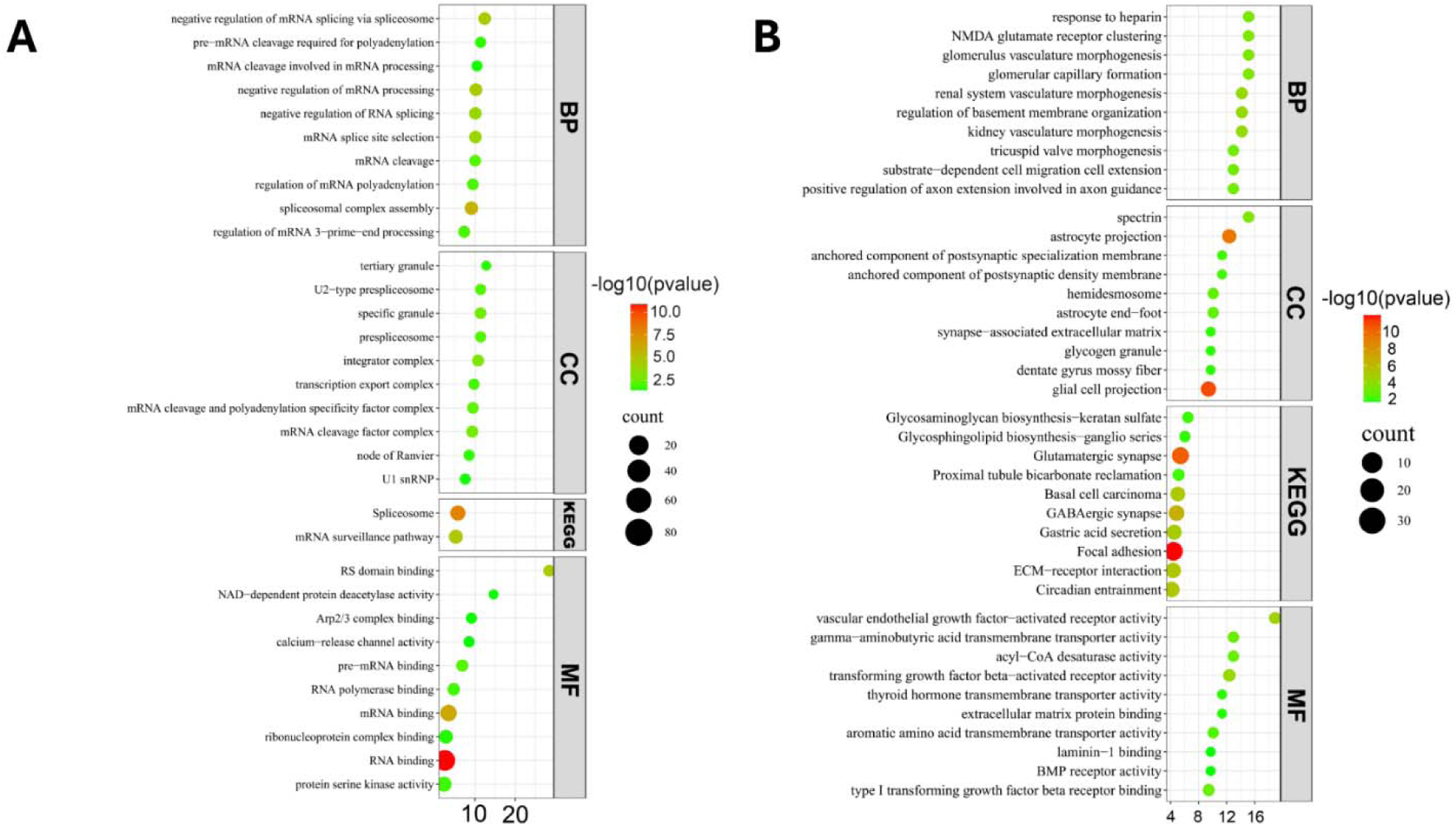
Pathway enrichment analyses for the differentially expressed genes identified in NPC1^mut^ female mouse brain samples. The dot plots depict pathways enriched by the (**A**) downregulated and (**B**) upregulated genes.

**Figure S6:**
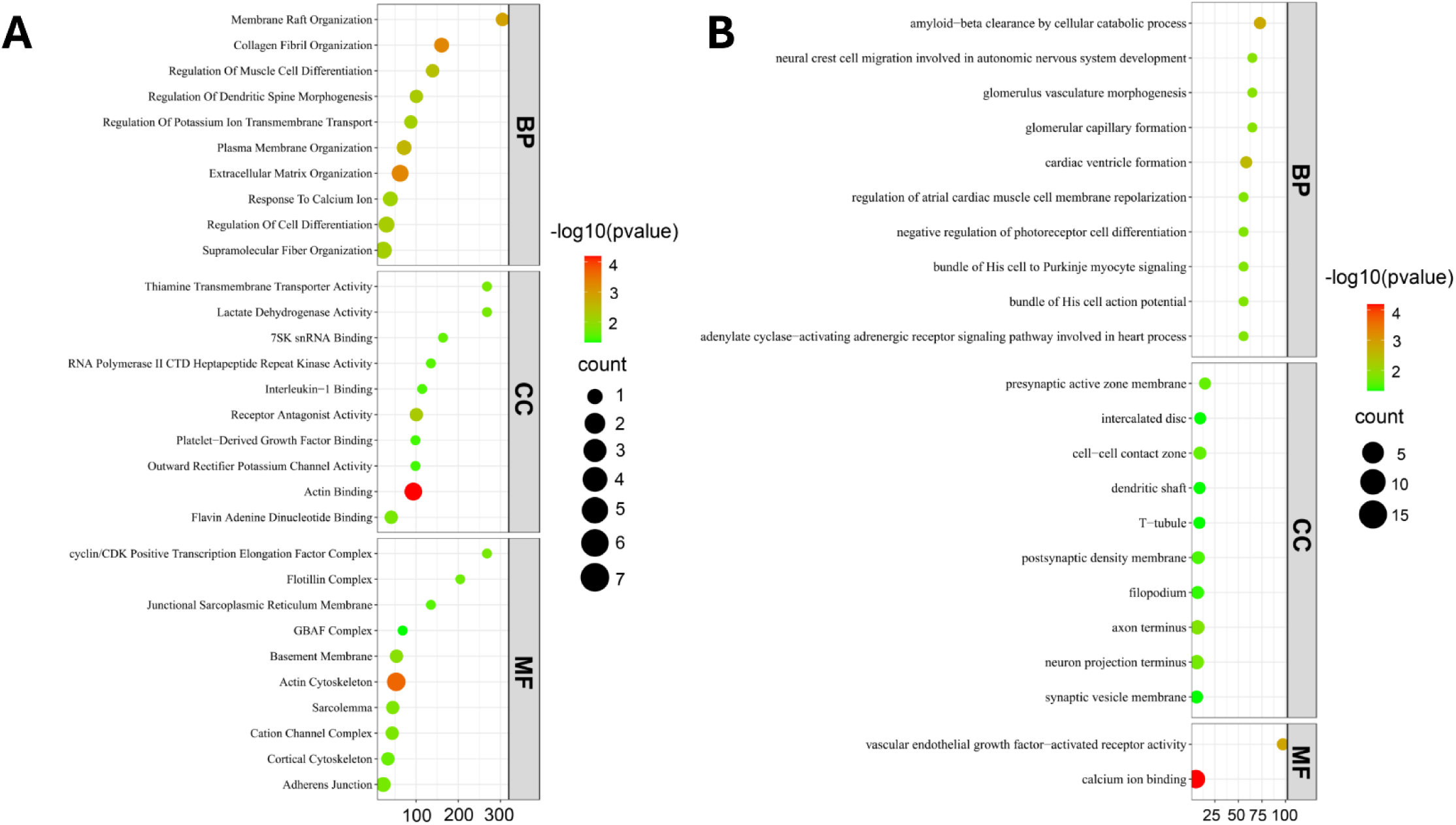
Pathway enrichment analyses for the differentially expressed genes identified in NPC1^mut^ male mouse brain samples. The dot plots depict pathways enriched by the (**A**) downregulated and (**B**) upregulated genes.

**Figure S7:**
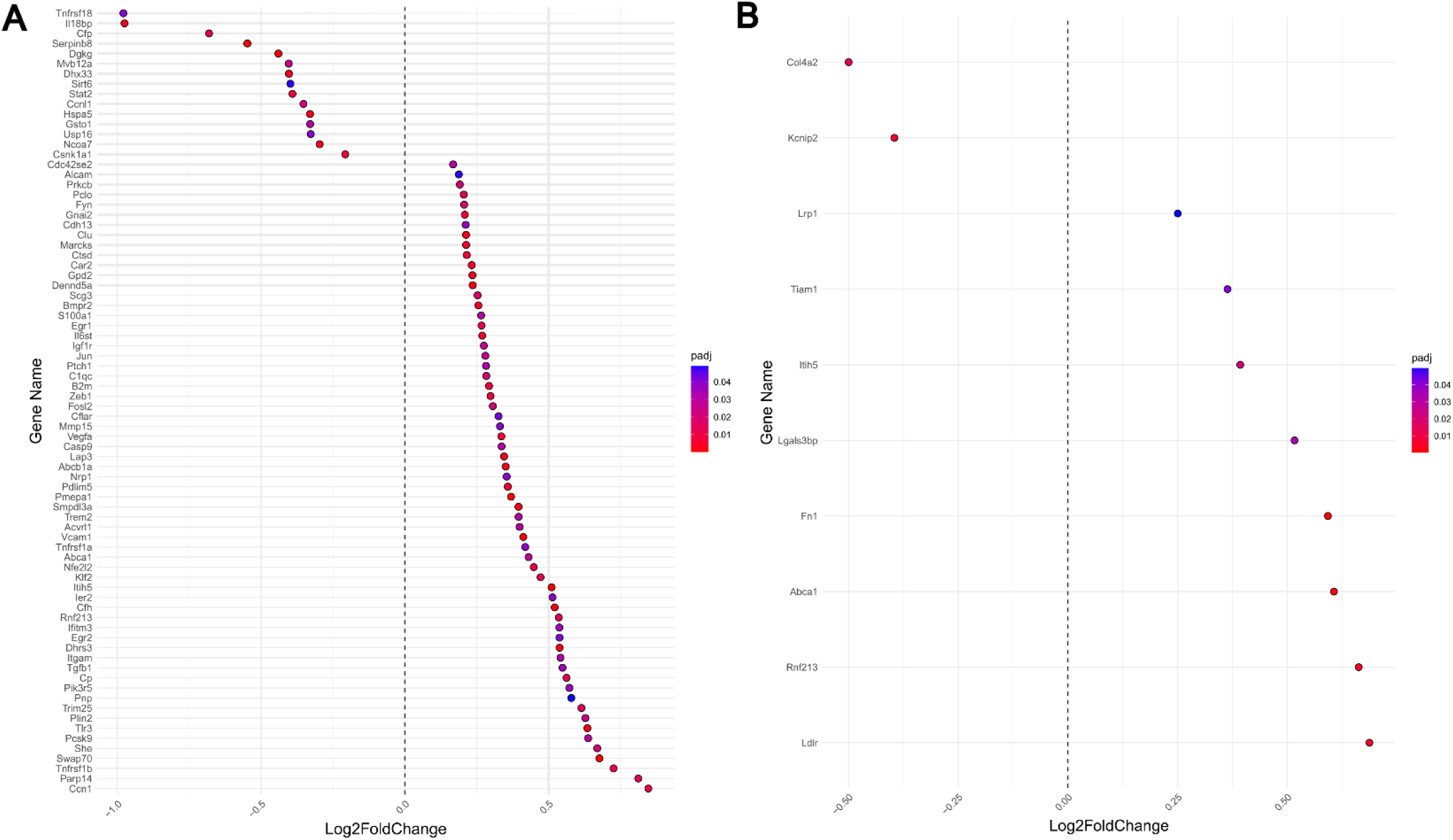
Altered expression of inflammatory genes in NPC1^mut^ mice. The dot plots depict the gene expression changes of inflammatory genes for (**A**) female and (**B**) male NPC1^mut^ mice.

**Figure S8:**
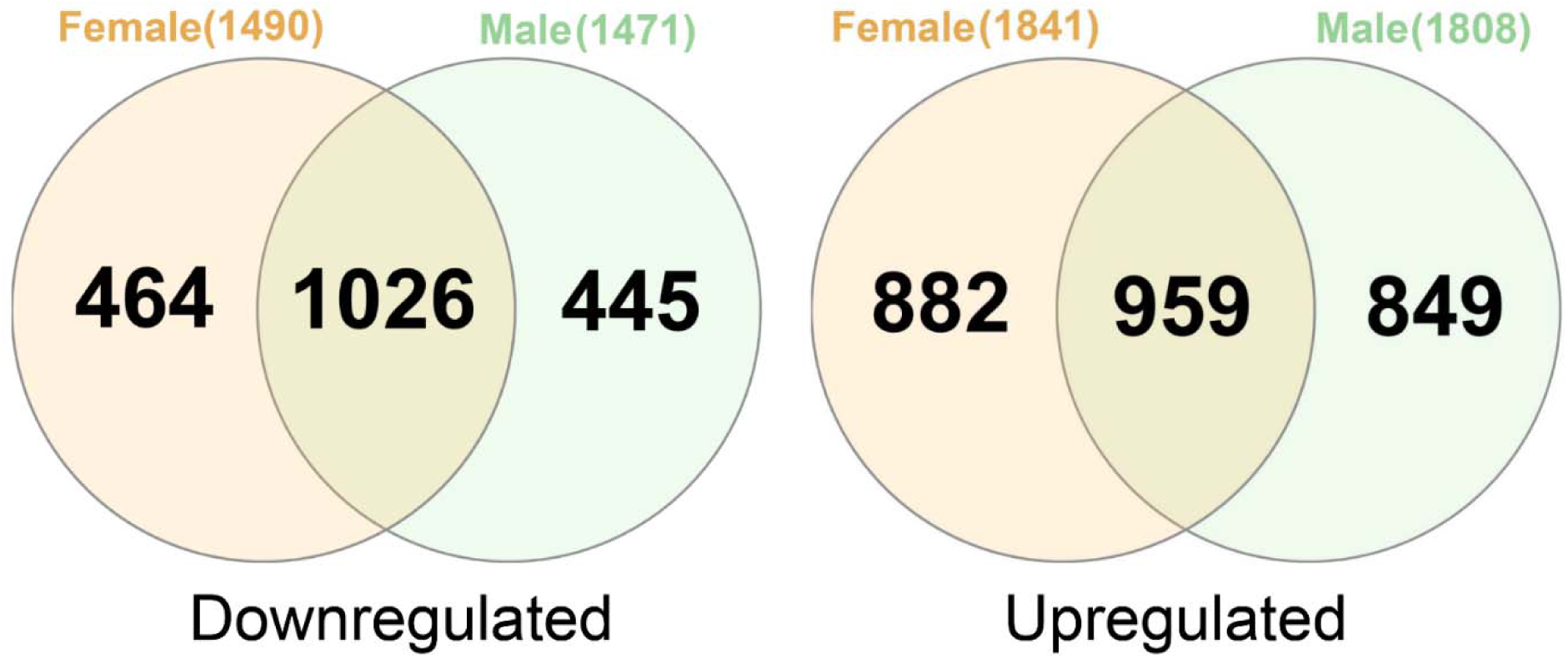
Venn diagrams depict the commonality analysis results on differentially expressed genes from female and male APP/PS1 mice relative to age and gender-matched WT controls. (**A)** There are 1,026 genes commonly downregulated and (**B**) 959 genes commonly upregulated between female and male frontal cortex samples of APP/PS1 mice. The full list of the common and unique DEGs between APP/PS1 female and male mice brain samples can be found in Supplementary File 4.

**Figure S9:**
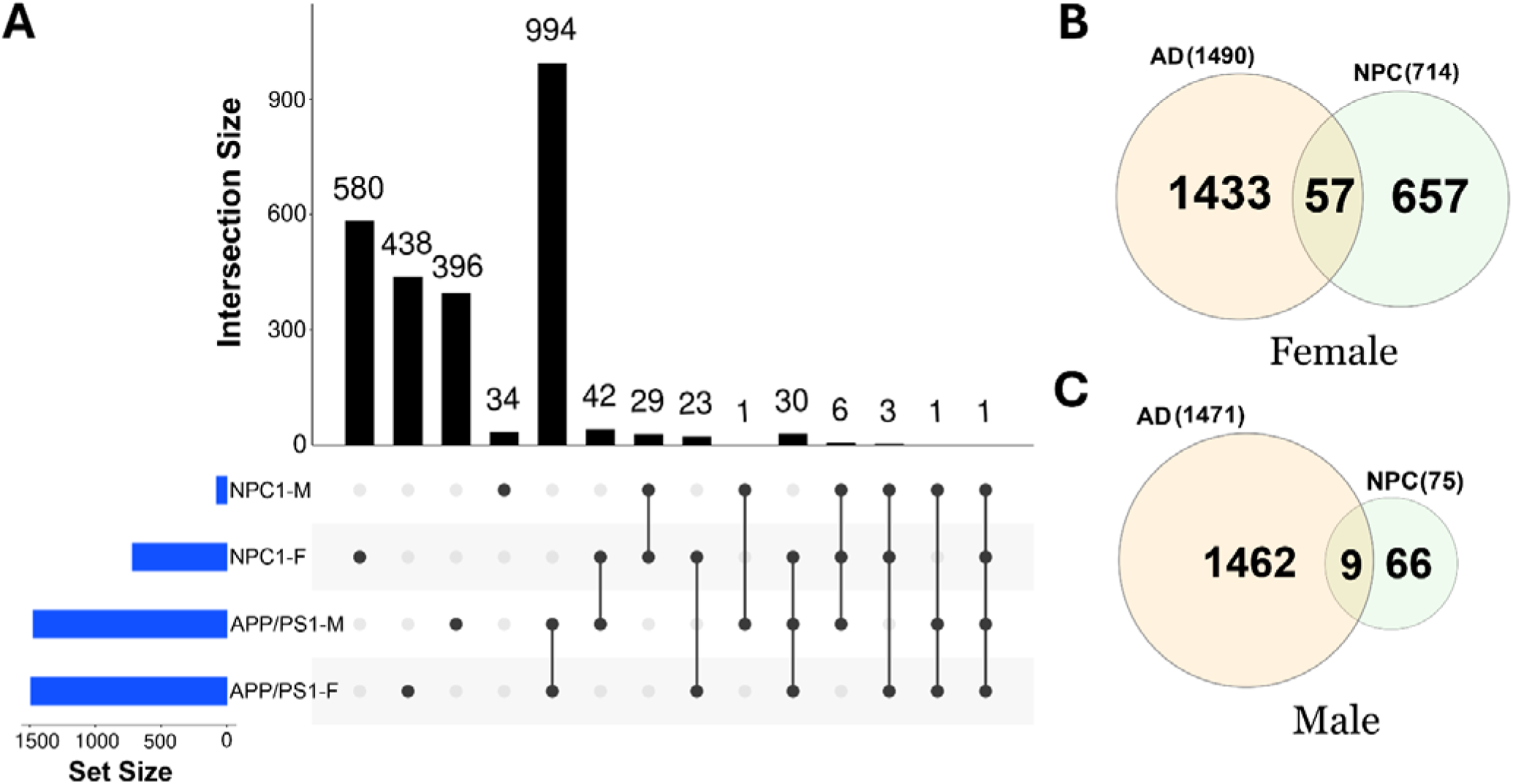
Cross-comparison of gene expression commonalities for the downregulated genes between frontal cortex samples of APP/PS1 and NPC1^mut^ mice. (**A**) Upset plot and (**B** and **C**) Venn diagrams depict the sex-specific downregulated genes. (**B**) There are 57 genes commonly downregulated between female NPC and AD mice. (**C**) There are 9 genes downregulated commonly between male NPC and AD mice. One gene, *MEG3,* is common to all four groups. The full list of the common and unique DEGs between APP/PS1 and NPC1^mut^ can be found in Supplementary File 5.

**Figure S10.**
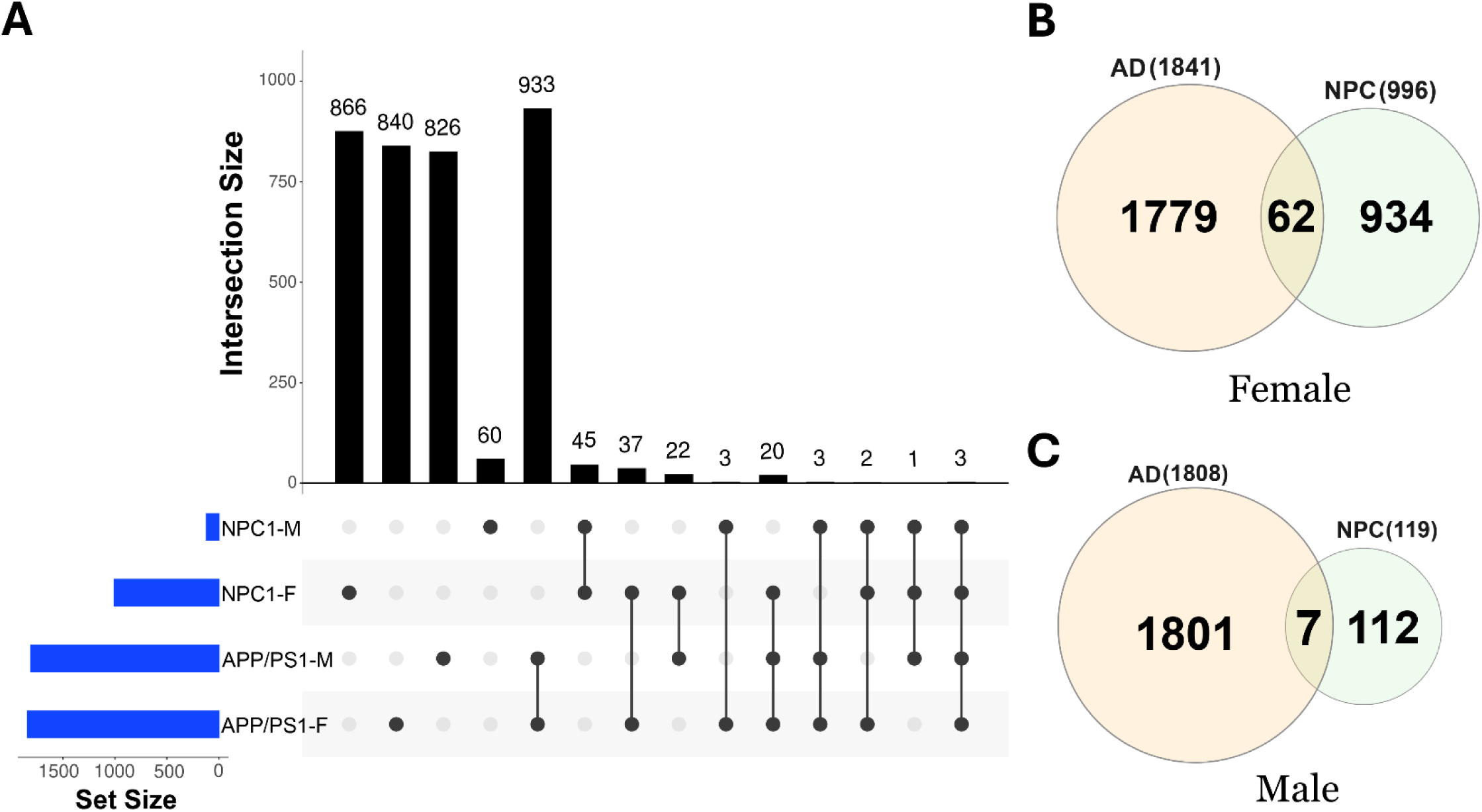
Cross-comparison of gene expression commonalities for the upregulated genes between frontal cortex samples of APP/PS1 and NPC1^mut^ mice. (**A**) Upset plot and (**B** and **C**) Venn diagrams depict the sex-specific upregulated genes. (**B**) There are 62 genes commonly upregulated between female NPC and AD mice. (**C**) Seven genes are commonly upregulated in male NPC and AD mice. Three genes, *PCDHB3*, *CAMK4*, and *TENM1*, are common to all four groups. The full list of the common and unique DEGs between APP/PS1 and NPC1^mut^ can be found in Supplementary File 5.

**Figure S11:**
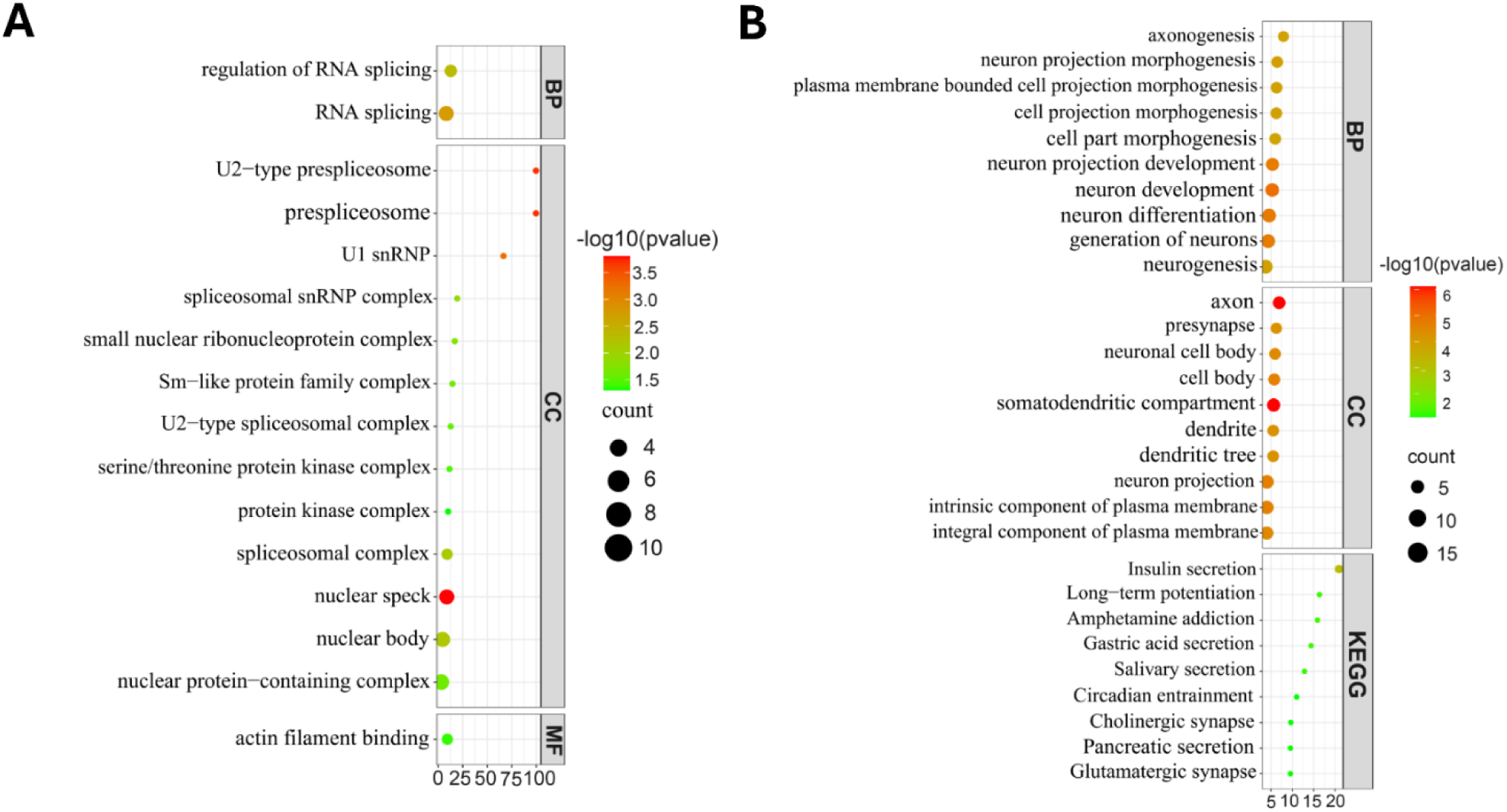
Pathway enrichment analyses for the commonly altered genes between female NPC and AD mice frontal cortex samples. Enrichment bubble plots depicting the downregulated (**A**) and upregulated (**B**) pathways associated with the differentially expressed genes common to Female NPC and AD mouse models. Among the downregulated pathways, many are associated with the spliceosome. However, the upregulated pathways are linked to neurogenesis, multiple synaptic responses, and neuronal differentiation. Similar analyses were not performed for the male NPC and AD mice due to their low number of shared genes.

**Figure S12:**
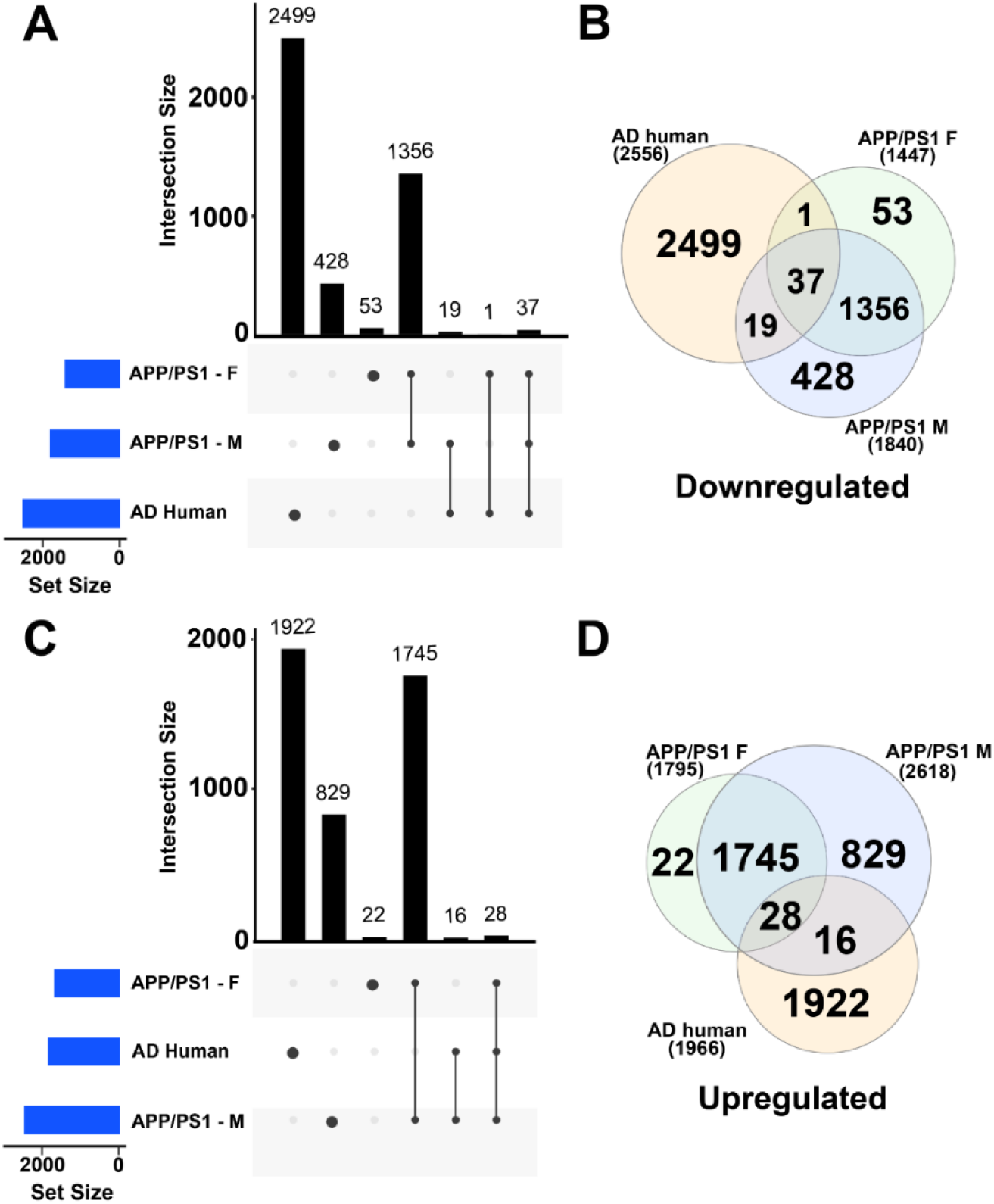
Cross-comparison of gene expression commonalities between frontal cortex samples of APP/PS1 mice and human AD brain samples. Upset plot (**A**) and (**B**) Venn diagram depicting the downregulated genes common to AD brain samples from humans and human orthologs from the male and female AD model of APP/PS1 mice. In total 37 genes are commonly downregulated in all groups. Only one downregulated gene, *HAVCR1*, is shared between AD brain and female AD mice frontal cortex samples. In contrast, 19 downregulated genes are common to AD brain and male AD mice samples. Upset plot (**C**) and (**D**) Venn diagram depicting the upregulated genes common to AD brain samples and human orthologs from the male and female AD mice. In total, 28 genes are commonly upregulated in all groups. There are no commonly upregulated genes between human AD brain and female AD mice samples. In contrast, 16 genes are commonly upregulated between AD brain and male AD mice samples. The complete list of the commonly altered genes can be found in Supplementary File 6.

## Supplementary Tables

**Table S1: Sample types and distribution of the number of altered gene expressions across different diseases and their commonalities.** Table depicting the parameters: differentially expressed genes per sample type, genes common across two or more diseases, and the p-value for Fisher’s exact test for randomness. Mosaic indicates analyses performed with a combination of human genes and human orthologs of mouse genes, several of which were identified to be common.

**Table S2:The list of all GEO accession numbers for the human data used in the study.** The columns correspond to the sample types, and the rows show the diseases.

**Table S3: Commonly regulated genes across sample types of different diseases.** The table lists commonly regulated genes, regulation types, molecular function, and disease and sample types in which they were identified.

**Table S4: Gene-miRNA interaction table.** A table containing important miRNA and genes found to be commonly altered across different diseases in our study and their disease associations identified by the number of publications.

## Supplementary Files

**File S1:** The full list of the top 50 positive and negative genes, along with their PCA scores for the organoid samples and the enriched pathways associated with their functions.

**File S2:** The complete lists of the DEGs for NPC1mut male and female mice brain samples.

**File S3:** Signature-based enriched functions of NPC1mut mice brain samples.

**File S4:** The list of the common and unique DEGs between APP/PS1 female and male mice brain samples.

**File S5:** The list of the common and unique DEGs in the brain samples of APP/PS1 and NPC1^mut^ (female and male) mice.

**File S6:** The list of the common DEGs in the brain samples of APP/PS, NPC1^mut^ mice and human AD.

**File S7:** GO enrichment terms for the genes commonly altered in NPC1^mut^ mice and human AD brain samples.

## References

1. Scheltens P, De Strooper B, Kivipelto M, Holstege H, Chételat G, Teunissen CE, et al. Alzheimer’s disease. The Lancet. 2021 Apr;397(10284):1577–90.

2. Chopade P, Chopade N, Zhao Z, Mitragotri S, Liao R, Chandran Suja V. Alzheimer’s and Parkinson’s disease therapies in the clinic. Vol. 8, Bioengineering and Translational Medicine. John Wiley and Sons Inc; 2023.

3. Fortea J, Quiroz YT, Ryan NS. Lessons from Down syndrome and autosomal dominant Alzheimer’s disease. Lancet Neurol. 2023 Jan;22(1):5–6.

4. Shimizu E, Goto-Hirano K, Motoi Y, Arai M, Hattori N. Symptoms and age of prodromal Alzheimer’s disease in Down syndrome: a systematic review and meta-analysis. Neurological Sciences. 2024 Jan 16;

5. Campos D, Monaga M. Mucopolysaccharidosis type I: current knowledge on its pathophysiological mechanisms. Metab Brain Dis. 2012 Jun 14;27(2):121–9.

6. Handen BL, Lott IT, Christian BT, Schupf N, OBryant S, Mapstone M, et al. The Alzheimer’s Biomarker Consortium-Down Syndrome: Rationale and methodology. Alzheimer’s & Dementia: Diagnosis, Assessment & Disease Monitoring. 2020 Jan 3;12(1).

7. Gomez W, Morales R, Maracaja-Coutinho V, Parra V, Nassif M. Down syndrome and Alzheimer’s disease: common molecular traits beyond the amyloid precursor protein. Aging. 2020 Jan 9;12(1):1011– 33.

8. Nixon RA. Niemann-Pick Type C Disease and Alzheimer’s Disease. Am J Pathol. 2004 Mar;164(3):757– 61.

9. Filocamo M, Tomanin R, Bertola F, Morrone A. Biochemical and molecular analysis in mucopolysaccharidoses: what a pediatrician must know. Ital J Pediatr. 2018 Nov 16;44(S2):129.

10. Handen BL. The Search for Biomarkers of Alzheimer’s Disease in Down Syndrome. Am J Intellect Dev Disabil. 2020 Mar 1;125(2):97–9.

11. Snow AD, Castillo GM. Specific proteoglycans as potential causative agents and relevant targets for therapeutic intervention in Alzheimer’s disease and other amyloidosis. Amyloid. 1997 Jan 6;4(2):135– 41.

12. Kobayashi H, Ariga M, Sato Y, Fujiwara M, Fukasawa N, Fukuda T, et al. P-Tau and Subunit c Mitochondrial ATP Synthase Accumulation in the Central Nervous System of a Woman with Hurler– Scheie Syndrome Treated with Enzyme Replacement Therapy for 12 Years. In 2018. p. 101–7.

13. Van Hoecke L, Van Cauwenberghe C, Dominko K, Van Imschoot G, Van Wonterghem E, Castelein J, et al. Involvement of the Choroid Plexus in the Pathogenesis of Niemann-Pick Disease Type C. Front Cell Neurosci. 2021 Oct 15;15.

14. Praggastis M, Tortelli B, Zhang J, Fujiwara H, Sidhu R, Chacko A, et al. A Murine Niemann-Pick C1 I1061T Knock-In Model Recapitulates the Pathological Features of the Most Prevalent Human Disease Allele. The Journal of Neuroscience. 2015 May 27;35(21):8091–106.

15. Jankowsky JL, Fadale DJ, Anderson J, Xu GM, Gonzales V, Jenkins NA, et al. Mutant presenilins specifically elevate the levels of the 42 residue β-amyloid peptide in vivo: evidence for augmentation of a 42-specific γ secretase. Hum Mol Genet. 2004 Jan 15;13(2):159–70.

16. Love MI, Huber W, Anders S. Moderated estimation of fold change and dispersion for RNA-seq data with DESeq2. Genome Biol. 2014 Dec 5;15(12):550.

17. Teichman G, Cohen D, Ganon O, Dunsky N, Shani S, Gingold H, et al. RNAlysis: analyze your RNA sequencing data without writing a single line of code. BMC Biol. 2023 Apr 7;21(1):74.

18. Heberle H, Meirelles GV, da Silva FR, Telles GP, Minghim R. InteractiVenn: a web-based tool for the analysis of sets through Venn diagrams. BMC Bioinformatics. 2015 May 22;16(1):169.

19. Zhou G, Soufan O, Ewald J, Hancock REW, Basu N, Xia J. NetworkAnalyst 3.0: a visual analytics platform for comprehensive gene expression profiling and meta-analysis. Nucleic Acids Res. 2019 Jul 2;47(W1):W234–41.

20. Shannon P, Markiel A, Ozier O, Baliga NS, Wang JT, Ramage D, et al. Cytoscape: A Software Environment for Integrated Models of Biomolecular Interaction Networks. Genome Res. 2003 Nov;13(11):2498–504.

21. Xie Z, Bailey A, Kuleshov M V., Clarke DJB, Evangelista JE, Jenkins SL, et al. Gene Set Knowledge Discovery with Enrichr. Curr Protoc. 2021 Mar 29;1(3).

22. Ge SX, Jung D, Yao R. ShinyGO: a graphical gene-set enrichment tool for animals and plants. Bioinformatics. 2020 Apr 15;36(8):2628–9.

23. Seal RL, Gordon SM, Lush MJ, Wright MW, Bruford EA. Genenames.org: The HGNC resources in 2011. Nucleic Acids Res. 2011 Jan;39(SUPPL. 1).

24. Eyre TA, Wright MW, Lush MJ, Bruford EA. HCOP: A searchable database of human orthology predictions. Brief Bioinform. 2007 Jan;8(1):2–5.

25. Wright MW, Eyre TA, Lush MJ, Povey S, Bruford EA. HCOP: The HGNC comparison of orthology predictions search tool. Mammalian Genome. 2005 Nov;16(11):827–8.

26. Võikar V, Rauvala H, Ikonen E. Cognitive deficit and development of motor impairment in a mouse model of Niemann-Pick type C disease. Behavioural Brain Research. 2002 Apr;132(1):1–10.

27. Tyshkovskiy A, Bozaykut P, Borodinova AA, Gerashchenko M V, Ables GP, Garratt M, et al. Identification and Application of Gene Expression Signatures Associated with Lifespan Extension. Cell Metab. 2019 Sep 3;30(3):573–593.e8.

28. Tyshkovskiy A, Ma S, Shindyapina A V, Tikhonov S, Lee SG, Bozaykut P, et al. Distinct longevity mechanisms across and within species and their association with aging. Cell. 2023 Jun 22;186(13):2929–2949.e20.

29. Robinson MD, McCarthy DJ, Smyth GK. edgeR: a Bioconductor package for differential expression analysis of digital gene expression data. Bioinformatics. 2010 Jan 1;26(1):139–40.

30. Subramanian A, Tamayo P, Mootha VK, Mukherjee S, Ebert BL, Gillette MA, et al. Gene set enrichment analysis: a knowledge-based approach for interpreting genome-wide expression profiles. Proc Natl Acad Sci U S A. 2005 Oct 25;102(43):15545–50.

31. McKay EC, Beck JS, Khoo SK, Dykema KJ, Cottingham SL, Winn ME, et al. Peri-Infarct Upregulation of the Oxytocin Receptor in Vascular Dementia. J Neuropathol Exp Neurol. 2019 May 1;78(5):436–52.

32. Lockstone HE, Harris LW, Swatton JE, Wayland MT, Holland AJ, Bahn S. Gene expression profiling in the adult Down syndrome brain. Genomics. 2007 Dec;90(6):647–60.

33. Kuehner JN, Chen J, Bruggeman EC, Wang F, Li Y, Xu C, et al. 5-hydroxymethylcytosine is dynamically regulated during forebrain organoid development and aberrantly altered in Alzheimer’s disease. Cell Rep. 2021 Apr;35(4):109042.

34. Czerminski JT, King OD, Lawrence JB. Large-scale organoid study suggests effects of trisomy 21 on early fetal neurodevelopment are more subtle than variability between isogenic lines and experiments. Front Neurosci. 2023 Feb 3;16.

35. Venkataraman L, Fair SR, McElroy CA, Hester ME, Fu H. Modeling neurodegenerative diseases with cerebral organoids and other three-dimensional culture systems: focus on Alzheimer’s disease. Stem Cell Rev Rep. 2022 Feb 12;18(2):696–717.

36. Lee SE, Shin N, Kook MG, Kong D, Kim NG, Choi SW, et al. Human iNSC-derived brain organoid model of lysosomal storage disorder in Niemann–Pick disease type C. Cell Death Dis. 2020 Dec 12;11(12):1059.

37. Papadimitriou C, Celikkaya H, Cosacak MI, Mashkaryan V, Bray L, Bhattarai P, et al. 3D Culture Method for Alzheimer’s Disease Modeling Reveals Interleukin-4 Rescues Aβ42-Induced Loss of Human Neural Stem Cell Plasticity. Dev Cell. 2018 Jul;46(1):85–101.e8.

38. Qiu J jun, Liu Y na, Wei H, Zeng F, Yan J bin. Single-cell RNA sequencing of neural stem cells derived from human trisomic iPSCs reveals the abnormalities during neural differentiation of Down syndrome. Front Mol Neurosci. 2023 Jun 15;16.

39. Swaroop M, Brooks MJ, Gieser L, Swaroop A, Zheng W. Patient iPSC-derived neural stem cells exhibit phenotypes in concordance with the clinical severity of mucopolysaccharidosis I. Hum Mol Genet. 2018 Oct 15;27(20):3612–26.

40. Leuba G, Vernay A, Vu D, Walzer C, Belloir B, Kraftsik R, et al. Differential expression of LMO4 protein in Alzheimer’s disease. Neuropathol Appl Neurobiol. 2004 Feb 27;30(1):57–69.

41. Yuan A, Rao M V., Sasaki T, Chen Y, Kumar A, Veeranna, et al. α-Internexin Is Structurally and Functionally Associated with the Neurofilament Triplet Proteins in the Mature CNS. The Journal of Neuroscience. 2006 Sep 27;26(39):10006–19.

42. Ching GY, Chien CL, Flores R, Liem RKH. Overexpression of α-Internexin Causes Abnormal Neurofilamentous Accumulations and Motor Coordination Deficits in Transgenic Mice. The Journal of Neuroscience. 1999 Apr 15;19(8):2974–86.

43. Cairns NJ, Zhukareva V, Uryu K, Zhang B, Bigio E, Mackenzie IRA, et al. α-Internexin Is Present in the Pathological Inclusions of Neuronal Intermediate Filament Inclusion Disease. Am J Pathol. 2004 Jun;164(6):2153–61.

44. Cairns NigelJ, Uryu K, Bigio EileenH, Mackenzie IanRA, Gearing M, Duyckaerts C, et al. α-Internexin aggregates are abundant in neuronal intermediate filament inclusion disease (NIFID) but rare in other neurodegenerative diseases. Acta Neuropathol. 2004 Sep 28;108(3).

45. Li QS, De Muynck L. Differentially expressed genes in Alzheimer’s disease highlighting the roles of microglia genes including OLR1 and astrocyte gene CDK2AP1. Brain Behav Immun Health. 2021 May;13:100227.

46. Wan YW, Al-Ouran R, Mangleburg CG, Perumal TM, Lee T V., Allison K, et al. Meta-Analysis of the Alzheimer’s Disease Human Brain Transcriptome and Functional Dissection in Mouse Models. Cell Rep. 2020 Jul;32(2):107908.

47. Chang JR, Ghafouri M, Mukerjee R, Bagashev A, Chabrashvili T, Sawaya BE. Role of p53 in Neurodegenerative Diseases. Neurodegener Dis. 2012;9(2):68–80.

48. Byrne JA, Tomasetto C, Garnier JM, Rouyer N, Mattei MG, Bellocq JP, et al. A screening method to identify genes commonly overexpressed in carcinomas and the identification of a novel complementary DNA sequence. Cancer Res. 1995 Jul 1;55(13):2896–903.

49. Byrne JA, Mattei MG, Basset P. Definition of the Tumor Protein D52 (TPD52) Gene Family through Cloning ofD52Homologues in Human (hD53) and Mouse (mD52). Genomics. 1996 Aug;35(3):523–32.

50. Shehata M, Bièche I, Boutros R, Weidenhofer J, Fanayan S, Spalding L, et al. Nonredundant Functions for Tumor Protein D52-Like Proteins Support Specific Targeting of TPD52. Clinical Cancer Research. 2008 Aug 15;14(16):5050–60.

51. Cao Q, Chen J, Zhu L, Liu Y, Zhou Z, Sha J, et al. A testis-specific and testis developmentally regulated tumor protein D52 (TPD52)-like protein TPD52L3/hD55 interacts with TPD52 family proteins. Biochem Biophys Res Commun. 2006 Jun;344(3):798–806.

52. Filippidis A, Carozza R, Rekate H. Aquaporins in Brain Edema and Neuropathological Conditions. Int J Mol Sci. 2016 Dec 28;18(1):55.

53. Hirt L, Price M, Benakis C, Badaut J. Aquaporins in neurological disorders. Clinical and Translational Neuroscience. 2018 Jan 7;2(1):2514183X1775290.

54. Moftakhar P, Lynch MD, Pomakian JL, Vinters H V. Aquaporin Expression in the Brains of Patients With or Without Cerebral Amyloid Angiopathy. J Neuropathol Exp Neurol. 2010 Dec;69(12):1201–9.

55. Barak M, Fedorova V, Pospisilova V, Raska J, Vochyanova S, Sedmik J, et al. Human iPSC-Derived Neural Models for Studying Alzheimer’s Disease: from Neural Stem Cells to Cerebral Organoids. Stem Cell Rev Rep. 2022 Feb 2;18(2):792–820.

56. Sorrentino F, Arighi A, Serpente M, Arosio B, Arcaro M, Visconte C, et al. Niemann-Pick Type C 1 (NPC1) and NPC2 Gene Variability in Demented Patients with Evidence of Brain Amyloid Deposition. Journal of Alzheimer’s Disease. 2021 Sep 28;83(3):1313–23.

57. Esposito M, Dubbioso R, Tozza S, Iodice R, Aiello M, Nicolai E, et al. In vivo evidence of cortical amyloid deposition in the adult form of Niemann Pick type C. Heliyon. 2019 Nov;5(11):e02776.

58. Mattsson N, Zetterberg H, Yanjanin NM, Månsson JE, Porter FD, Blennow K. Increased cerebrospinal fluid amyloid β levels in Niemann-Pick disease patients: Implications for Alzheimer’s disease. Alzheimer’s & Dementia. 2010 Jul;6(4S_Part_8).

59. Nixon RA. Niemann-Pick Type C Disease and Alzheimer’s Disease. Am J Pathol. 2004 Mar;164(3):757– 61.

60. Colombo A, Dinkel L, Müller SA, Sebastian Monasor L, Schifferer M, Cantuti-Castelvetri L, et al. Loss of NPC1 enhances phagocytic uptake and impairs lipid trafficking in microglia. Nat Commun. 2021 Feb 24;12(1):1158.

61. Rudnitskaya EA, Kozlova TA, Burnyasheva AO, Kolosova NG, Stefanova NA. Alterations of hippocampal neurogenesis during development of Alzheimer’s disease-like pathology in OXYS rats. Exp Gerontol. 2019 Jan;115:32–45.

62. Pfeffer A, Munder T, Schreyer S, Klein C, Rasińska J, Winter Y, et al. Behavioral and psychological symptoms of dementia (BPSD) and impaired cognition reflect unsuccessful neuronal compensation in the pre-plaque stage and serve as early markers for Alzheimer’s disease in the APP23 mouse model. Behavioural Brain Research. 2018 Jul;347:300–13.

63. Rudnitskaya EA, Kolosova NG, Stefanova NA. Impact of changes in neurotrophic supplementation on development of Alzheimer’s disease-like pathology in OXYS rats. Biochemistry (Moscow). 2017 Mar 11;82(3):318–29.

64. Mehla J, Lacoursiere SG, Lapointe V, McNaughton BL, Sutherland RJ, McDonald RJ, et al. Age-dependent behavioral and biochemical characterization of single APP knock-in mouse (APPNL-G-F/NL-G-F) model of Alzheimer’s disease. Neurobiol Aging. 2019 Mar;75:25–37.

65. Radbruch H, Mothes R, Bremer D, Seifert S, Köhler R, Pohlan J, et al. Analyzing Nicotinamide Adenine Dinucleotide Phosphate Oxidase Activation in Aging and Vascular Amyloid Pathology. Front Immunol. 2017 Jul 31;8.

66. Keck BJ, Lakoski JM. N-Ethoxycarbonyl-2-ethoxy-1,2-dihydroquinoline (EEDQ) administration for studies of 5-HT1A receptor binding site inactivation and turnover. Brain Research Protocols. 1997 Oct;1(4):364–70.

67. López-Toledano MA, Shelanski ML. Increased Neurogenesis in Young Transgenic Mice Overexpressing Human APPSw,Ind. Journal of Alzheimer’s Disease. 2007 Nov 19;12(3):229–40.

68. Murer MG, Boissiere F, Yan Q, Hunot S, Villares J, Faucheux B, et al. An immunohistochemical study of the distribution of brain-derived neurotrophic factor in the adult human brain, with particular reference to Alzheimer’s disease. Neuroscience. 1999 Feb;88(4):1015–32.

69. Jiang Q, Shan K, Qun-Wang X, Zhou RM, Yang H, Liu C, et al. Long non-coding RNA-MIAT promotes neurovascular remodeling in the eye and brain. Oncotarget. 2016 Aug 2;7(31):49688–98.

70. Sun C, Huang L, Li Z, Leng K, Xu Y, Jiang X, et al. Long non-coding RNA MIAT in development and disease: a new player in an old game. J Biomed Sci. 2018 Dec 13;25(1):23.

71. Fasolo F, Jin H, Winski G, Chernogubova E, Pauli J, Winter H, et al. Long Noncoding RNA MIAT Controls Advanced Atherosclerotic Lesion Formation and Plaque Destabilization. Circulation. 2021 Nov 9;144(19):1567–83.

72. Qi D, Hou X, Jin C, Chen X, Pan C, Fu H, et al. HNSC exosome-derived MIAT improves cognitive disorders in rats with vascular dementia via the miR-34b-5p/CALB1 axis. Am J Transl Res. 2021;13(9):10075–93.

73. Li J, Ye J. Chronic intermittent hypoxia induces cognitive impairment in Alzheimer’s disease mouse model via postsynaptic mechanisms. Sleep and Breathing. 2024 Jan 24;

74. Qureshi YH, Patel VM, Berman DE, Kothiya MJ, Neufeld JL, Vardarajan B, et al. An Alzheimer’s Disease-Linked Loss-of-Function CLN5 Variant Impairs Cathepsin D Maturation, Consistent with a Retromer Trafficking Defect. Mol Cell Biol. 2018 Oct 1;38(20).

75. Rossi C, Distaso M, Raggi F, Kusmic C, Faita F, Solini A. Lacking P2X7-receptors protects substantia nigra dopaminergic neurons and hippocampal-related cognitive performance from the deleterious effects of high-fat diet exposure in adult male mice. Front Nutr. 2024 Jan 25;11.

76. Qin Z, Zhou X, Gomez-Smith M, Pandey NR, Lee KFH, Lagace DC, et al. LIM Domain Only 4 (LMO4) Regulates Calcium-Induced Calcium Release and Synaptic Plasticity in the Hippocampus. The Journal of Neuroscience. 2012 Mar 21;32(12):4271–83.

77. Qiu B, Li X, Sun X, Wang Y, Jing Z, Zhang X, et al. Overexpression of aquaporin-1 aggravates hippocampal damage in mouse traumatic brain injury models. Mol Med Rep. 2014 Mar;9(3):916–22.

78. McNair K, Spike R, Guilding C, Prendergast GC, Stone TW, Cobb SR, et al. A Role for RhoB in Synaptic Plasticity and the Regulation of Neuronal Morphology. The Journal of Neuroscience. 2010 Mar 3;30(9):3508–17.

79. Kamasani U, Prendergast GC. Genetic response to DNA damage: Proapoptotic targets of RhoB include modules for p53 response and susceptibility to alzheimer’s disease. Cancer Biol Ther. 2005 Mar 27;4(3):282–8.

80. Teplova M, Hafner M, Teplov D, Essig K, Tuschl T, Patel DJ. Structure–function studies of STAR family Quaking proteins bound to their in vivo RNA target sites. Genes Dev. 2013 Apr 15;27(8):928–40.

81. Sakers K, Liu Y, Llaci L, Lee SM, Vasek MJ, Rieger MA, et al. Loss of Quaking RNA binding protein disrupts the expression of genes associated with astrocyte maturation in mouse brain. Nat Commun. 2021 Mar 9;12(1):1537.

82. Fernandez-Martos CM, King AE, Atkinson RAK, Woodhouse A, Vickers JC. Neurofilament light gene deletion exacerbates amyloid, dystrophic neurite, and synaptic pathology in the APP/PS1 transgenic model of Alzheimer’s disease. Neurobiol Aging. 2015 Oct;36(10):2757–67.

83. Vega FM, Ridley AJ. The RhoB small GTPase in physiology and disease. Small GTPases. 2018 Sep 3;9(5):384–93.

84. Aguilar BJ, Zhu Y, Lu Q. Rho GTPases as therapeutic targets in Alzheimer’s disease. Alzheimers Res Ther. 2017 Dec 15;9(1):97.

85. Brabeck C, Beschorner R, Conrad S, Mittelbronn M, Bekure K, Meyermann R, et al. Lesional Expression of RhoA and RhoB following Traumatic Brain Injury in Humans. J Neurotrauma. 2004 Jun;21(6):697–706.

86. Baek JH, Schmidt E, Viceconte N, Strandgren C, Pernold K, Richard TJC, et al. Expression of progerin in aging mouse brains reveals structural nuclear abnormalities without detectible significant alterations in gene expression, hippocampal stem cells or behavior. Hum Mol Genet. 2015 Mar 1;24(5):1305–21.

87. Lee BC, Péterfi Z, Hoffmann FW, Moore RE, Kaya A, Avanesov A, et al. MsrB1 and MICALs regulate actin assembly and macrophage function via reversible stereoselective methionine oxidation. Mol Cell. 2013 Aug 8;51(3):397–404.

88. P A H, Kommaddi RP, Ravindranath V. Differential expression of Mical2 and its activator (Sema3a) in Alzheimer’s disease mouse model. Alzheimer’s & Dementia. 2022 Dec 20;18(S3).

