## Supplemental Table 1 for "A disease similarity approach identifies short-lived Niemann-Pick type C disease mice with accelerated brain aging as a novel mouse model for Alzheimer’s disease and aging research"

**Table S1: Sample types and distribution of the number of altered gene expressions across different diseases and their commonalities.** Table depicting the parameters: differentially expressed genes per sample type, genes common across two or more diseases, and the p-value for Fisher’s exact test for randomness. Mosaic indicates analyses performed with a combination of human genes and human orthologs of mouse genes, several of which were identified to be common.

|  |  |  | **Human** | | | | **Mouse** | |  |  |
| --- | --- | --- | --- | --- | --- | --- | --- | --- | --- | --- |
|  | **Expression** | **Sample type** | **AD** | **DS** | **NPC** | **MPS I** | **APP/PS1** | **Npc1** | **Common** | **p-value** |
| **Human** | **Downregulated** | **Brain samples** | 2556 | 202 | X | X | X | X | 96 (AD - DS) | 6.02E-06 |
|  |  | **Organoids** | 1053 | 792 | 474 | X | X | X | 2 (AD - DS - NPC), 102 (AD - DS), 19 (AD - NPC) | 4.99E-06 |
|  |  | **iNSC** | 33 | 7889 | X | 596 | X | X | 10 (AD - DS), 2 (AD - MPS I), 3 (AD - DS - MPS I) | 4.99E-07 |
|  | **Upregulated** | **Brain samples** | 1966 | 476 | X | X | X | X | 117 (AD - DS) | 4.72E-06 |
|  |  | **Organoids** | 191 | 948 | 783 | X | X | X | 4 (AD - DS), 6 (AD - NPC), 1 (AD - DS - NPC) | 4.99E-06 |
|  |  | **iNSC** | 156 | 470 | X | 1010 | X | X | 8 (AD - DS), 10 (AD - MPS I), 2 (AD - DS - MPS I) | 4.99E-07 |
| **Mouse** | **Downregulated** | **Brain samples - female** | X | X | X | X | 1490 | 714 | 57 (APP/PS1 - Npc1) | 4.62E-05 |
|  |  | **Brain samples - male** | X | X | X | X | 1471 | 75 | 9 (APP/PS1 - Npc1) | 6.50E-04 |
|  | **Upregulated** | **Brain samples - female** | X | X | X | X | 1841 | 996 | 62 (APP/PS1 - Npc1) | 3.50E-04 |
|  |  | **Brain samples - male** | X | X | X | X | 1808 | 119 | 7 (APP/PS1 - Npc1) | 5.06E-04 |
| **Mosaic** | **Downregulated** | **Brain samples** | 2556 | X | X | X | X | 714 (F) | 68 (AD - Npc1_f), 6 (AD - Npc1_m), 27 (Npc1_f, Npc1_m), 6 (AD - Npc1_f - Npc1_m) | 2.20E-07 |
|  |  |  |  | X | X | X | X | 75 (M) |  |  |
|  | **Upregulated** | **Brain samples** | 1966 | X | X | X | X | 996 (F) | 119 (AD - Npc1_f), 6 (AD - Npc1_m), 44 (Npc1_f - Npc1_m), 6 (AD - Npc1_f - Npc1_m) | 2.20E-16 |
|  |  |  |  | X | X | X | X | 119 (M) |  |  |
|  | **Downregulated** | **Brain samples** | 2556 | X | X | X | 1490 (F) | X | 1 (AD - h - APP/PS1 - f), 19 (AD - h - APP/PS1 - m), 1356 (APP/PS1 - f - APP/PS1 - m), 37 (AD - h - APP/PS1 - f - APP/PS1 - m) | 2.20E-16 |
|  |  |  |  | X | X | X | 1471 (M) | X |  |  |
|  | **Upregulated** | **Brain samples** | 1966 | X | X | X | 1841 (F) | X | 16 (AD - h - APP/PS1 - m), 1745 (APP/PS1 - f - APP/PS1 - m), 28 (AD - h - APP/PS1 - f - APP/PS1 - m) | 3.47E-170 |
|  |  |  |  | X | X | X | 1808 (M) | X |  |  |
