## Supplemental Table 2 for "A disease similarity approach identifies short-lived Niemann-Pick type C disease mice with accelerated brain aging as a novel mouse model for Alzheimer’s disease and aging research"

**Table S2**: The list of all GEO accession numbers for the human data used in the study. The columns correspond to the sample types, and the rows show the diseases.

| **Disease** | **Brain samples** | **Organoids** | **iNSCs** |
| --- | --- | --- | --- |
| **AD** | **GSE122063**Click or tap here to enter text. | **GSE151818**Click or tap here to enter text. | **GSE78117**Click or tap here to enter text. |
|  | **Age range**: Control [Female (73-87), Male (60-91)], AD [Female (63-89), Male (79-91)]  **Number of samples**: Control (11) (M/F:5/6), AD (12) (M/F:3/9) | **Sample number**: Control (9), AD (3) | **Sample number**: Control (3), AD (2) |
| **DS** | **GSE5390**Click or tap here to enter text. | **GSE222365**Click or tap here to enter text. | **GSE208625**Click or tap here to enter text. |
|  | **Age range**: Control [Female (61)], DS [Female (61, 76), Male (63)] **Number of samples**: Control (1) (M/F:0/1), DS (3) (M/F:1/2) | **Sample number**: Control (4), DS (4) | **Sample number**: Control (3), DS (3) |
| **NPC** | --- | **GSE157676**Click or tap here to enter text. | --- |
|  |  | **Sample number**: Control (3), NPC (3) |  |
| **MPS I** | --- | --- | **GSE111906**Click or tap here to enter text. |
|  |  |  | **Sample number**: Control (3), MPS I HS (3), MPS I H (3), MPS I S (3) |
