## Supplemental Table 3 for "A disease similarity approach identifies short-lived Niemann-Pick type C disease mice with accelerated brain aging as a novel mouse model for Alzheimer’s disease and aging research"

**Table S3**: **Commonly regulated genes across sample types of different diseases.** The table lists commonly regulated genes, regulation types, molecular function, and disease and sample types in which they were identified.

| **Gene** | **Regulation** | **Function** | **AD** | **DS** | **NPC** | **MPS I** | **Sample type** |
| --- | --- | --- | --- | --- | --- | --- | --- |
| *QKI* | Down | Myelinization, RNA binding | X | X | X | -- | Organoids |
| *RHOB* | Down | GTPase activity, apoptosis after DNA damage | X | X | X | -- | Organoids |
| *PTPRO* | Down | Tyrosine phosphatase activity | X | -- | -- | X | Brain samples, iNSCs |
| *NETO2* | Down | Affects Synaptic transmission by slowing EPSC decay | X | X | -- | X | Brain samples, Organoids, iNSCs |
| *SHISA2* | Down | FGF and WNT signaling | X | X | -- | -- | Organoids, iNSCs |
| *SPON2* | Down | Innate immune response initiation, cell adhesion | X | -- | -- | -- | Brain samples, iNSCs |
| *INA* | Down | Neuron morphogenesis, intracellular transport to axons and dendrites | X | X | X | -- | Brain samples, Organoids |
| *LMO4* | Down | Transcription cofactor negatively modulates interneuron genes in motor neurons | X | X | X | -- | Brain samples, Organoids |
| *AQP1* | Up | Obligate water channel, involved in CSF production | X | X | -- | X | Brain samples, iNSCs |
| *TPD52L1* | Up | Calcium signaling, positive regulation of MAP3K5-induced apoptosis | X | X | X | -- | Brain samples, Organoids |
| *ASNS* | Up | Asparagine biosynthesis | X | -- | -- | -- | iNSCs |
| *DDIT3* | Up | CCAAT/enhancer-binding transcription factor | X | -- | -- | -- | iNSCs |
| *TNMD* | Up | Associated with age-related macular degeneration, angiogenesis inhibitor | X | -- | -- | -- | iNSCs |
| *ZNF248* | Up | DNA-binding transcription repressor | X | X | -- | -- | Brain samples |
| *PCDH1* | Up | Neural cell adhesion and cell-cell interaction processes | X | X | -- | -- | Brain samples, iNSCs |
| *STON2* | Up | Synaptic vesicle recycling, neurotransmission | X | X | -- | -- | Brain samples, iNSCs |
| *GDF15* | Up | pleiotropic cytokine, stress response to hypoxia | X | X | -- | X | Brain samples, iNSCs |
