## Supplemental Table 4 for "A disease similarity approach identifies short-lived Niemann-Pick type C disease mice with accelerated brain aging as a novel mouse model for Alzheimer’s disease and aging research"

**Table S4**: Table containing important miRNA, genes interacting with them, and their disease associations depicting the number of studies.

| **miRNA (hsa-mir)** | **Genes** | **AD** | **DS** | **NPC** | **MPS I** |
| --- | --- | --- | --- | --- | --- |
| 29a-3p | AQP1, TPD52L1, SFRP | 1 | -- | -- | -- |
| 9-5p | INA, LMO4, GDF15 | 1 | -- | -- | -- |
| 125a-3p | CLSTN2 | 1 | -- | -- | -- |
| 125b-5p | NETO2 | 1 | -- | -- | -- |
| 1-3p | GDF15, NETO2, AQP1, TPD52L1 | 2 | -- | -- | -- |
| 101-3p | NETO2, INA, LMO4 | 2 | -- | -- | -- |
| 155-5p | INA, LMO4, AQP1, TPD52L1, NETO2, GDF15 | 3 | 1 | -- | -- |
| 128-3p | NETO2, CLSTN2, GDF15 | 4 | -- | -- | -- |
| 16-5p | INA, LMO4, AQP1, TPD52L1, NETO2 | 9 | -- | -- | -- |
| 124-3p | GDF15, NETO2 | 19 | 3 | -- | -- |
| 107 | CLSTN2 | 20 | -- | -- | -- |
| 7b-5p | AQP1, TPD52L1 | -- | -- | -- | -- |
| 145-5p | NETO2, CLSTN2 | -- | -- | -- | -- |
| 7-5p | CLSTN2 | -- | -- | -- | -- |
| 133a-3p | NETO2, GDF15 | -- | -- | -- | -- |
| 10b-5p | SFRP2 | -- | -- | -- | -- |
| 218-5p | SFRP2 | -- | -- | -- | -- |
| 429 | NETO2, GDF15, ADM2 | -- | -- | -- | -- |
| 23b-3p | GDF15 | -- | -- | -- | -- |
